# TRACR: an anterograde transneuronal tracing system for genetic access across synapses and longitudinal circuit analysis

**DOI:** 10.64898/2026.02.08.704659

**Authors:** Mostafa Ibrahimi, Madison T. Gray, Pierre-Luc Rochon, Sofia M. Eiras, Joseph J. Lee, Paula Pietraszkiewicz, Sara Dabiri, Janice L. Hicks, Michael W. Salter, Marc Boisvert, Marie-Ève Paquet, Sheena A. Josselyn, Kirill A. Martemyanov, Arjun Krishnaswamy, Valerie A. Wallace, Julie L. Lefebvre

## Abstract

Following neural signals as they converge onto and diverge from individual neurons is central to understanding circuit function and disease-related dysfunction. Existing anterograde transneuronal tracers are limited by cytotoxicity and incomplete genetic control over connected partners. To address these limitations, we adapted synthetic Notch designs to create TRanssynaptic Anterograde Circuit Readout (TRACR). Binding of the engineered ligand-receptor across synapses induces TRE-driven reporter transcription, enabling characterization of postsynaptic neurons. TRACR provides segregated genetic access to pre- and postsynaptic populations, and can be combined with markers, sensors, or effectors to expand circuit analysis. By applying TRACR at multiple synapses in the mouse visual system, we show that TRACR labels postsynaptic partners of sensory neurons, long-range projections and local inhibitory interneurons. TRACR signaling is reversible and fails to activate when synapses are absent or disrupted. Together, TRACR is an accessible, AAV-deliverable transneuronal reporting tool for longitudinal analysis of circuit assembly, degeneration, and repair.

**HIGHLIGHTS:**

- TRACR adapts the synNotch system to signal across synapses for tracing postsynaptic targets.
- TRACR identifies local and long-range postsynaptic targets in the mouse visual system.
- TRACR activation requires intact synaptic connectivity rather than proximity.
- TRACR signals are reversible, diminishing upon synapse loss and activating following assembly.

## INTRODUCTION

A central goal of neuroscience is to explain how neural circuits encode information, and how they change in disease states. Achieving this goal requires knowing which cell types are synaptically connected within circuits, yet this remains a major technical challenge. Functional approaches, such as electrophysiology or optogenetic recordings, can reveal connectivity between individual neurons or defined populations, but are difficult to integrate with circuit-wide anatomical analyses. Structural approaches, including immunohistochemistry and electron microscopy, can identify synapse location and synaptic abnormalities, but not which neuronal cell types are connected within an intact circuit. Transneuronal tracing methods that label across connected neurons and leverage existing molecular neuroscience tools offer a powerful strategy for mapping neuronal connectivity *in vivo*. An ideal circuit-mapping tool would identify the connected cell types in the direction of information flow, work across diverse types of synapses in local and long-range circuits, and allow genetic access to both partners for molecular profiling and functional manipulations. As circuits remodel during development and disease, a tracing strategy designed to capture connectivity changes over time would also open the door to studying circuit assembly, degeneration, and repair.

Current viral and intercellular tracing strategies can link neural identity to connectivity, but their performance varies across synapse types, pathways, and experimental timelines. In mammals, the synapse crossing property of neurotropic viruses has been leveraged for transsynaptic tracing. The glycoprotein-deleted rabies virus is the most widely used strategy for retrograde monosynaptic tracing, enabling transgene transfer from defined starter neurons to mark, monitor or manipulate their presynaptic inputs (Wickersham, Finke, et al. 2007; Wickersham, Lyon, et al. 2007; Osakada and Callaway 2013). However, their rapid cytotoxicity typically limits the duration of analyses, although recent modifications may extend their utility for longer-term studies (Chatterjee et al. 2018; Jin et al. 2024). Anterograde viral tools, including engineered HSV, VSV, YFV-17D and AAV1-based systems, can label postsynaptic partners from a given neural type but are often limited by undesired retrograde spread, incomplete genetic access to starter and/or target cells, cytotoxicity, and biosafety or regulatory constraints (Lo and Anderson 2011; Beier et al. 2013; Zingg et al. 2017; Zingg et al. 2020; Li et al. 2021; Xiong et al. 2022; Bouin et al. 2024; Jin et al. 2024). Non-viral strategies that exploit intercellular transfer or neurotransmission are promising (e.g., WGA, ATLAS, BAcTrace) but are limited to certain synapse types and their utility varies across circuits (Cachero et al. 2020; Tsai et al. 2022; Rivera et al. 2025). Importantly, most existing approaches—viral and non-viral—yield largely static snapshots via material transfer or permanent reporter induction, limiting the ability to resolve circuit remodeling. Together, these limitations highlight the need for a genetically encoded strategy that reports on synaptic contact without relying on viral spread, while providing reversible readouts of connectivity.

To address these gaps, we developed a transneuronal tracing system based on synthetic Notch signaling (synNotch) that converts synaptic contact into a customizable transcriptional readout. SynNotch receptors retain the core Notch regulatory and transmembrane domains but replace the extracellular domain with a single-chain variable fragment or nanobody that binds a user-defined ligand (e.g. GFP), and the intracellular domain with a transcriptional activator that drives a chosen output gene (Morsut et al. 2016). Ligand binding triggers intramembrane proteolysis by endogenous proteases, releasing the transcriptional activator to enter the nucleus and activate reporter expression. SynNotch has been used *in vitro* to report cell–cell contacts and program cellular behaviors (Morsut et al. 2016; Roybal et al. 2016; Toda et al. 2018; Choe et al. 2021; Malaguti et al. 2022; Reddy et al. 2024), and *in vivo* to trace contacts during mouse vascular development (Zhang et al. 2022). In *Drosophila*, the related TRACT system demonstrated that synNotch-based contact sensing can detect anterograde synaptic partners in select circuits (Huang et al. 2016; Huang et al. 2017). Together, these advances establish synNotch as a promising platform for genetically encoded transsynaptic tracing in mammals.

Here, we present TRACR - *Transsynaptic Anterograde Circuit Readout* – an AAV-deliverable synNotch system for identifying postsynaptic partners of genetically-defined neurons *in vivo*. TRACR uses a tetracycline-controlled transactivator (tTA)-dependent transcriptional readout to label postsynaptic partners, while preserving segregated access to pre- and postsynaptic populations for the expression of markers, sensors or effectors. Through a series of experiments in the mouse visual system, we show that TRACR can be restricted to genetically defined starter cells, and used to efficiently and specifically label postsynaptic partners across both local and long-range connections. By evaluating the parameters of TRACR signalling at the photoreceptor-bipolar synapse and in mouse models in which these synapses fail to form or subsequently degenerate, we show that reporter induction requires intact synaptic connectivity rather than simply proximity of synaptic terminals. Moreover, TRACR signals diminish following presynaptic degeneration and report synaptic reassembly, demonstrating its reversibility and utility for tracing changes in connectivity. Together, we present a modular TRACR toolkit for transsynaptic tracing that can be broadly adopted to study circuits of interest in the mouse nervous system, as well circuit remodeling in development and disease.

## RESULTS

### TRACR system design

To develop a ligand-receptor based tracing method for mammalian circuits, we adapted the synthetic Notch (synNotch) system to convert synaptic contact into a reporter signal that can be visualized in vivo. Building on canonical synNotch designs (Morsut et al. 2016), we designed a three-component Sender-Receiver-Reporter system that signals as follows (Figure 1A): ‘Sender’ cells display a membrane-tethered GFP ligand at presynaptic terminals. ‘Receiver’ neurons express a chimeric synNotch receptor comprising an extracellular anti-GFP nanobody (LaG17), the Notch core regulatory and transmembrane domains, and a cleavable intracellular transcriptional module, tTA. Ligand binding exerts mechanical force across the Notch core, triggering the endogenous proteolytic cascade, and liberating the tTA to translocate to the nucleus and drive expression of a Reporter transgene under tTA-responsive elements (TRE). We designed the components to be delivered via adeno-associated virus (AAV), targeting Sender expression to genetically defined presynaptic populations using Cre/LoxP strategies or cell type-specific promoters, and enabling identification of target cells among ubiquitous Receiver and Reporter-expressing neurons (Figure 1B).

**Figure 1:**
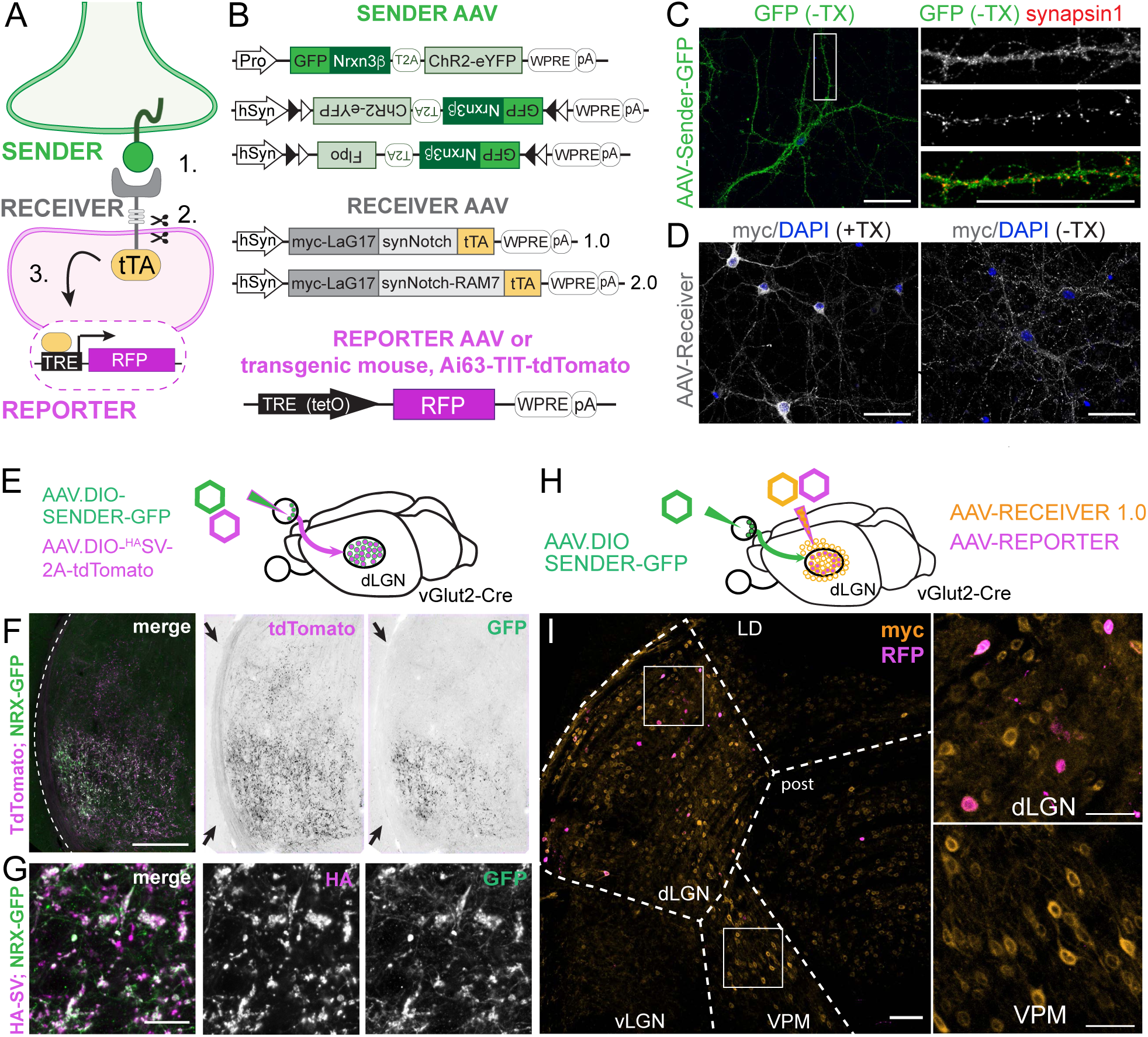
TRACR system and transneuronal tracing of long-range connections. **(A)** Schematic of TRACR. A presynaptic Sender neuron (green) displays an NRX–GFP ligand enriched at terminals. A postsynaptic Receiver neuron (magenta) expresses synNotch receptors comprising an extracellular GFP nanobody (LaG17), the Notch core regulatory and transmembrane domains (grey), and the intracellular transcriptional activator tTA. Ligand binding across the synapse [1] triggers intramembrane proteolysis of synNotch [2] and release of tTA, which translocates to the nucleus to activate a TRE-responsive Reporter [3], labeling the postsynaptic neuron with RFP. **(B)** TRACR components. Sender AAVs encode NRX-GFP, a 2A peptide, and a marker or effector transgene. Sender expression is driven by either a cell-type-specific promoter (Pro) or a pan-neuronal promoter (hSyn), and can be Cre-dependent (DIO, double-floxed inverted orientation). Receiver AAVs encode two synNotch-tTA variants (1.0 or 2.0) with an N-terminal myc. TRE-driven reporters used here include AAV.TRE-mRuby2 and Ai63 (TIT-tdTomato) mice. **(C)** Sender NRX-GFP localization in cultured DIV14 cortical neurons co-infected with AAV.hSyn-Cre and AAV.hSyn-DIO-NRXGFP-T2A-Flpo. Surface staining (green; no Triton X-100 (-TX)) shows NRX–GFP at the plasma membrane and at presynaptic sites overlapping with Synapsin1 (red, with sequential permeabilization; inset enlarged at right). **(D)** synNotch localization in cultured neurons infected with Receiver AAV.hSyn-mycLaG17-synNotch-tTA. Immunostaining for myc (white) shows synNotch along neurites and at the cell surface (right, no Triton X-100). **(E)** Schematic of strategy to visualize Sender NRX-GFP within presynaptic terminals of retinal ganglion cells (RGCs) in dorsal lateral geniculate nuclei (dLGN). AAVs encoding Sender (hSyn-DIO-NRXGFP-T2A-Flpo) and HA-tagged synaptic vesicle (SV) marker (hSyn-DIO-9XHA-SV-2A-tdTomato) were delivered to retinas of *vGluT2-Cre* mice. **(F)** Representative confocal image of dLGN (dashed outline) shows NRX-GFP (green) overlap with RGC terminal axons (tdTomato from SV marker, magenta). Inverted images show the enrichment of GFP (right) along with tdTomato-positive terminal arborizations (right), and the relative absence of GFP along axon tracts (middle, arrows). **(G)** Localization of NRX-GFP (green) within RGC terminal boutons and overlapping with HA-tagged SV marker (magenta). **(H)** AAV strategy to express Sender (hSyn-DIO-NRXGFP-T2A-ChR2-YFP) in the retina, and Receiver (hSyn-mycLaG17-synNotch-tTA) and Reporter (TRE-mRuby2) in the thalamus of *vGluT2-Cre* mice. Mice were injected at 3-4 postnatal weeks and harvested at 2 months of age. **(I)** Image of thalamus showing TRACR labeled retinothalamic connections. Receiver neurons (myc, orange) are broadly distributed across thalamic nuclei. Reporter expression (RFP, magenta) is restricted to myc-positive Receiver cells within the target region (dLGN) but absent from non-target nuclei, including laterodorsal (LD) and ventral posteromedial (VPM) nuclei. **Right:** enlarged insets. Scale bars: 50 µm (C,D); 200 µm (F, I); 20µm (F, bottom); and 10 µm (I, right).

A critical challenge in adapting synNotch for transneuronal tracing is to confine contact-dependent signaling to synaptic interfaces. After evaluating previous synapse-targeting strategies (Kim et al. 2011; Yamagata and Sanes 2012; Martell et al. 2016; Coomer et al. 2025), we tethered the GFP ligand to the transmembrane and cytoplasmic domains of Neurexin-3β (Figure 1B). These Neurexin-3β domains are highly similar to those in Neurexin-1β, and promote targeting to axon terminals and surface membrane insertion (Fairless et al. 2008; Gokce and Südhof 2013; Klatt et al. 2021). To minimize artifacts, we omitted the extracellular domains of Neurexin-3β which can drive ectopic synaptogenic interactions when overexpressed (Kim et al. 2011; Tsetsenis et al. 2014). To augment cellular processing of the Nrx3β (NRX)-GFP ligand, we incorporated signal peptide and membrane trafficking sequences shown to improve surface delivery of engineered opsin proteins (Zhao et al. 2008; Gradinaru et al. 2010).

The Sender AAV constructs encoding the NRX-GFP ligand include a promoter of choice or a Cre-dependent design to restrict expression to genetically identified presynaptic neuronal populations (Figure 1B). A modular bicistronic design enables co-expression of NRX–GFP with fluorescent markers, recombinases or effectors to support downstream investigations such as anatomical tracing and optogenetic interrogations (Figure 1B). In cultured neurons, co-transfection of an AAV Sender (AAV.hSyn-DIO-NRXGFP-T2A-FlpO) along with AAV.hSyn-Cre produced robust expression and trafficking of NRX–GFP throughout neurites (Figure 1C). Consistent with surface presentation, staining extracellular GFP under non-permeabilized conditions labeled NRX–GFP puncta that overlap with presynaptic marker, synapsin-1 (Figure 1C).

For unbiased tracing of postsynaptic cells, we created Receiver plasmids to express synNotch ubiquitously in neuronal cells without post-synaptic tethering (Figure 1B). This choice was intended to minimize interference with synNotch activation, and to permit detection across diverse excitatory and inhibitory synapse types, which share few proteins at the postsynaptic density, including neuroligins (Bemben et al. 2015). The tTA (Tet-OFF) was selected as the transcriptional effector based on its prior demonstrated performance in synNotch assays and the extensive implementation of tTA-TRE transgenic platforms in the mouse nervous system (Madisen et al. 2015; Morsut et al. 2016; Malaguti et al. 2022; Zhang et al. 2022). Transfection of Receiver-AAV in cultured neurons confirmed robust surface expression of the myc-tagged LaG17-synNotch-tTA fusion proteins along neuronal processes (Figure 1D). For tTA-dependent reporter expression, we used available AAVs (Chan et al. 2017) and transgenic mice expressing red fluorescent protein (RFP) from the TRE promoter (Ai63-TIT-tdT, (Daigle et al. 2018) (Figure 1B). To minimize the ligand-independent activation reported in contexts of high synNotch expression, we constructed a second Receiver (Receiver 2.0, Figure 1B) with a hydrophobic protein sequence inserted in the C-terminus of the Notch core (RAM7, (Yang et al. 2020)).

We designated this entire system as TRACR – *<u>Tr</u>*anssynaptic *<u>A</u>*nterograde *<u>C</u>*ircuit *<u>R</u>*eadout. We posited that in TRACR, NRX–GFP on presynaptic terminals binds synNotch on postsynaptic partners to release tTA and drive Reporter expression, resulting in RFP labeling of synaptic targets. We next tested the sensitivity and specificity of TRACR signaling across multiple synapses in the mouse visual system.

### TRACR identifies long-range connections

We first sought to assess the TRACR system in vivo in the retinothalamic projections. The mouse retinothalamic projection has well-defined anatomical boundaries, where retinal ganglion cells (RGCs) send axons to the thalamus and most prominently innvervate the dorsal lateral geniculate nucleus (dLGN). We predicted that TRACR signaling will induce reporter expression in dLGN neurons, but not in the surrounding thalamic nuclei that lack RGC inputs, such as the ventral posteromedial nucleus (VPM) (Martersteck et al. 2017).

To visualize the NRX-GFP ligand within RGC terminals in the dLGN, we co-injected Cre-dependent AAVs encoding the Sender and an HA-tagged presynaptic marker (Chantranupong et al. 2020) into retinas of vGluT2-Cre mice, which label almost all RGCs (Martersteck et al. 2017) (Figure 1E). GFP positive axon terminals were distributed throughout known retinal projection fields and overlapped with the HA-synaptophysin, confirming a presynaptic localisation of the Sender-GFP ligand (Figure 1F, G).

We next delivered the AAVs encoding the Receiver (Receiver 1.0) and TRE-mRuby Reporter via stereotaxic injection to the thalamus, encompassing the dLGN and surrounding nuclei. The myc-tagged Receiver was broadly expressed across the injected thalamic area, including the dLGN, VPM, posterior complex, and lateral dorsal nucleus (Figure 1H, I). To control for ligand-independent activation and reporter leakiness, we injected Receiver and Reporter AAVs at different dilutions, and without the Sender virus. While delivery of high titer of the Receiver and Reporter AAVs caused leaky RFP signals, delivery of increasingly diluted AAVs revealed graded decrease in reporter signals to absent or rare RFP-positive cells (Supp. Figure 1). Thus, titrating dosages of the Receiver and Reporter AAVs in the absence of Sender AAV in the region of interest is required for non-leaky tracing, similar to titrating helper and Reporter AAVs for retrograde tracing using rabies virus (Lavin et al. 2020).

When all TRACR components were delivered, RFP expression induced by synNotch signaling in putative postsynaptic partners was almost exclusively restricted to the dLGN (Figure 1I). The RFP labeling within the dLGN, coupled with its conspicuous absence in the adjacent Receiver-expressing but non-synaptically-connected nuclei, demonstrates that TRACR identifies targets within a long-range anatomical projection.

### TRACR identifies interneuron targets in local circuitry

We next tested TRACR to dissect local neuronal connections, a task that remains challenging with established transsynaptic tracers. To evaluate TRACR’s resolution and specificity in a dense short-range network, we turned to the inner plexiform layer (IPL) of the retina (Figure 2A). The IPL is an ideal test site where neurites of over 100 molecularly defined neuron types stratify into specific sublaminae to establish distinct functional circuits (Shekhar et al. 2016; Tran et al. 2019; Yan et al. 2020). Focusing on a well-known circuit for direction selectivity, we tested the specificity of TRACR for tracing a local circuit by evaluating the: i) morphology of Reporter-positive neurons, including overlap with pre-synaptic neurites, ii) expression of appropriate molecular markers, and iii) functional analysis.

**Figure 2:**
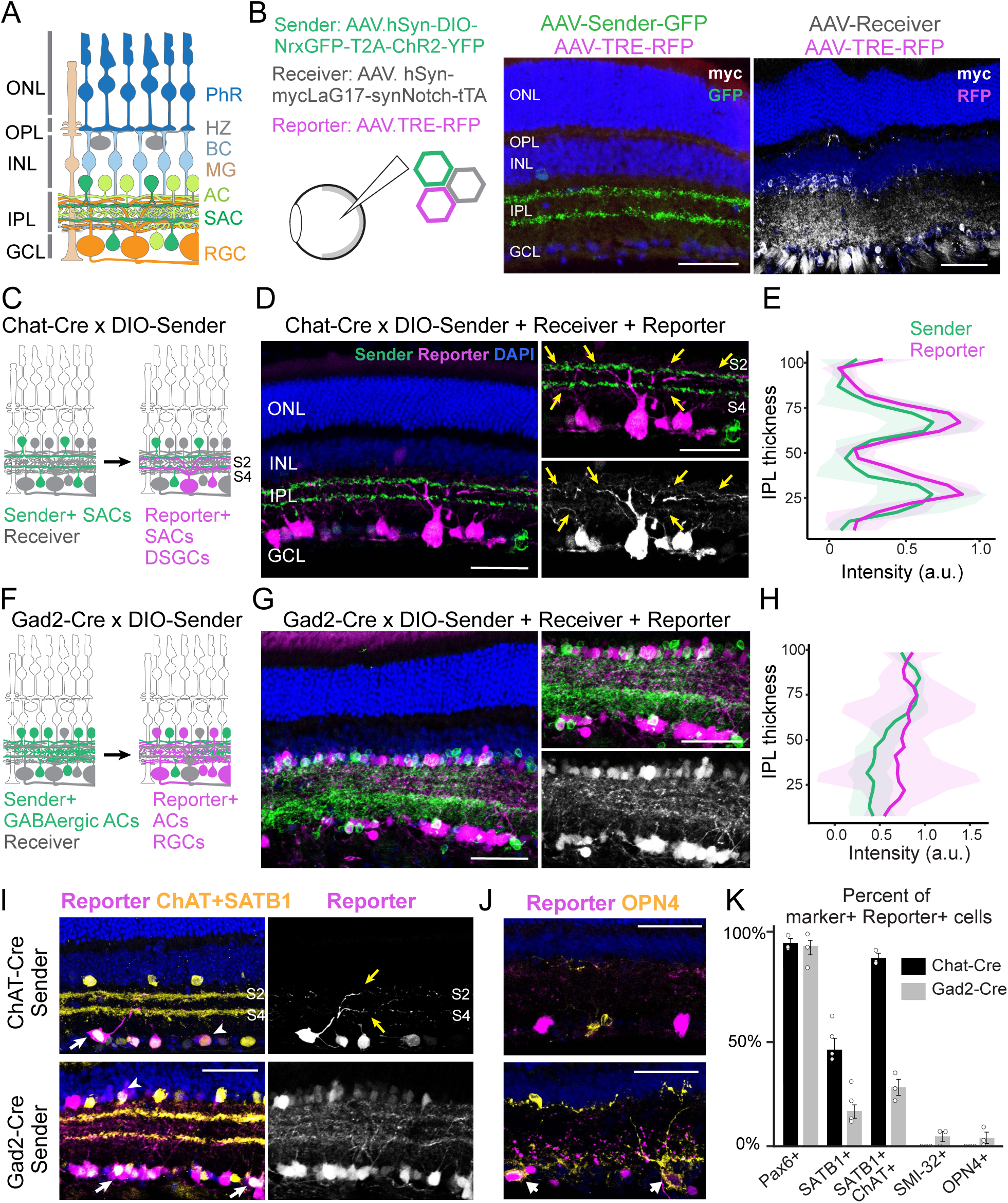
TRACR identifies interneuron targets in the inner retina. **(A)** Schematic of the retina. Photoreceptors (PhR) in the outer nuclear layer (ONL) synapse in the outer plexiform layer (OPL) onto bipolar cell (BP) and horizontal cell (HZ) dendrites. BPs and amacrine cells (ACs) project to the inner plexiform layer (IPL), where they form stereotyped, lamina-specific connections with retinal ganglion cell (RGC) subtypes. Starburst amacrine cells (SACs, green) are inhibitory ACs that stratify in two IPL sublaminae and synapse onto other SACs and direction-selective RGCs (DSGCs). **(B) Left:** AAV TRACR components and controls: Sender, Receiver, and TRE-driven Reporter. Mice received AAV intravitreal injections at 3-4 postnatal weeks and harvested at 2 months of age. **Middle:** *ChAT-Cre* retina injected with Receiver and Reporter AAVs but lacking Sender shows Receiver expression (myc, white) without TRE-driven RFP (middle). **Right:** Retinas injected with Sender and Reporter AAVs shows GFP ligand (green) but no RFP induction. **(C)** Schematic of TRACR strategy to label SAC targets. In *ChAT-Cre* retinas, Sender (green) is restricted to SACs (green), whereas Receiver and Reporter are broadly expressed across inner retinal neurons (grey). Reporter induction (magenta) is predicted in SACs and DSGCs. **(D)** Representative image of TRACR-labeled *ChAT-Cre* retina. Sender-GFP (green) and Reporter-positive neurites (RFP, magenta) co-stratify in the expected IPL sublaminae. Insets highlight overlapping neurites (yellow arrowheads). **(E)** Quantification of Sender-positive and Reporter-positive neurite stratification in *ChAT-Cre* retinas. Line profiles of GFP and RFP fluorescence across the IPL show strong anatomical correspondence (cos θ similarity index = 0.89 ± 0.05). Data are shown as mean (dark line) ± SEM (light ribbon); n = 3 images per retina, 4 retinas from 4 animals. **(F)** TRACR strategy to label GABAergic AC targets. In *Gad2-Cre* retinas, Sender (GFP, green) is expressed across broad AC populations. Reporter induction (RFP, magenta) is predicted in diverse ACs and RGCs. **(G)** Representative section of TRACR-labeled *Gad2-Cre* retina. Sender-positive (green) and Reporter-positive neurites (magenta) are throughout the IPL. **(H)** Line profiles of GFP and RFP-positive neurites in *Gad2-Cre* retinas show overlap (cos θ similarity index = 0.93 ± 0.07). RFP signals show distinct distributions from *ChAT-Cre* retinas in E (cos θ similarity index = 0.60 ± 0.02, p < 0.01). Data are shown as mean (dark line) ± SEM (light ribbon); n = 3 images per retina, 4 retinas from 4 animals. Similarity indices differ significantly from E, p < 0.01, one-way ANOVA. **(I)** Molecular validation of TRACR targets in *ChAT-Cre* retinas. **Left:** Reporter-labeled cells (magenta) include SACs (ChAT, yellow; arrowheads) and DSGCs (Satb1, yellow; arrows). Middle: RFP-positive cell shows bistratified dendrite morphology to S2/S4 typical of ON-OFF DSGC (center, yellow arrows). **Right:** Reporter cells exclude non-target RGC types such as melanopsin-positive ipRGCs (Opn4, yellow). **(J)** Molecular validation of TRACR targets in *Gad2-Cre* retinas using markers as in (I). **(K)** TRACR targets labeled by *ChAT-Cre*-driven Senders differ from *Gad2-Cre*-driven Senders. Plots show the percentage of marker-positive Reporter-positive neurons for Pax6 (pan ACs and RGCs), SATB1 (ON–OFF DSGCs), SATB1+ChAT (DSGCs/SACs), SMI-32 (α-RGCs), and Opn4 (ipRGCs). 3–5 retinas per condition. p = 0.64 (Pax6), 0.034 (SATB1), 0.0015 (SATB1+ChAT), 0.046 (SMI-32), 0.037 (Opn4). Scale bars: 50 µm (B, D, G, I, J).

We evaluated TRACR in retinas of ChAT-Cre mice, which express Cre exclusively in starburst amacrine cells (SACs) (Lefebvre et al. 2012) (Figure 2B,C). SACs are GABAergic interneurons that shape the movement responses of direction-selective retinal ganglion cells (DSGCs). Their dendrites stratify in two IPL sublaminae (s2 and s4), where they synapse primarily onto monostratified ON and bistratified ON–OFF DSGCs, and onto other SACs (Yoshida et al. 2001; Briggman et al. 2011; Yonehara et al. 2011; Chen et al. 2016; Ding et al. 2016; Brombas et al. 2017; Mauss et al. 2017), with additional RGC types receiving weaker innervation (Beier et al. 2013; Baden et al. 2016; Wang and Zhang 2023). We predicted that TRACR Reporter activation will occur mainly in SACs and DSGCs, with Sender and RFP-positive neurites co-stratifying in s2 and s4 (Figure 2C).

TRACR Reporter induction required all three components: co-injection of Receiver and Reporter AAVs without Sender, or Sender and Reporter AAVs without Receiver, yielded minimal RFP (Figure 2B). When all TRACR AAVs are delivered in ChAT-Cre retinas, Reporter-positive neurons were observed in both the inner nuclear layer (INL) and ganglion cell layer (GCL), with processes overlapping with Sender neurites in S2 and S4 in the IPL (cosine similarity index = 0.89± 0.05, Figure 2D, E). To compare the SAC-driven TRACR output to other presynaptic neurons, we performed parallel experiments in Gad2-Cre mice, which label a broader class of GABAergic amacrine cells (Figure 2F). As expected, Sender neurite lamination in Gad2-Cre retinas was broader than observed in ChAT-Cre retinas, but these neurites also overlapped with Reporter labelled neurites (Figure 2G, H). Importantly, Reporter signals in Gad2-Cre retinas were significantly different than that of ChAT-Cre (cosine similarity index = 0.60 ± 0.02, *p* <0.01 vs ChAT-Cre and p=0.01 vs Gad2-Cre), confirming that TRACR Reporter signal is dependent on the Sender population.

Molecular characterization of TRACR signals supported circuit specificity: 90% of Reporter-positive neurons in ChAT-Cre retinas co-expressed either ChAT (SACs) or SATB1 (ON–OFF DSGCs), markers for SAC target cells (189/211 cells, n = 5 retinas) (Lefebvre et al. 2012; Peng et al. 2017) (Figure 2I-K). In contrast, only 16% (33/209 cells, n = 3 retinas) of Reporter-positive neurons in Gad2-Cre retinas expressed markers of SACs or ON-OFF DSGCs. Consistent with known wiring, no reporter co-localization was detected (0/126 and 0/101 cells, respectively; n = 3 retinas) with SMI-32 (α-RGCs) or Opn4 (ipRGCs) in ChAT-Cre retinas, two RGC types not receiving SAC input (Viney et al. 2007; Baden et al. 2016) (Figure 2J,K). As expected, Gad2-Cre TRACR labeling included these cell types at frequencies consistent with their population abundance (α-RGCs, 5.1%; ipRGCs, 6.0%) (Hattar et al. 2002; Tran et al. 2019).

We next performed loose-patch recordings from Reporter-positive and -negative RGCs in ChAT-Cre retinal explants treated with TRACR (Figure 3A, B). RFP-positive RGCs (n = 32 cells) showed predominately ON-OFF responses to a full field flash, however some ON- or OFF-RGCs were encountered (Figure 3C). This was in contrast with random sampling of nearby RFP-negative RGCs (n = 42 cells) which tended to be predominantly ON- or OFF-RGCs. These results suggest that Reporter-positive RGCs are enriched for ON-OFF DSGCs.

**Figure 3.**
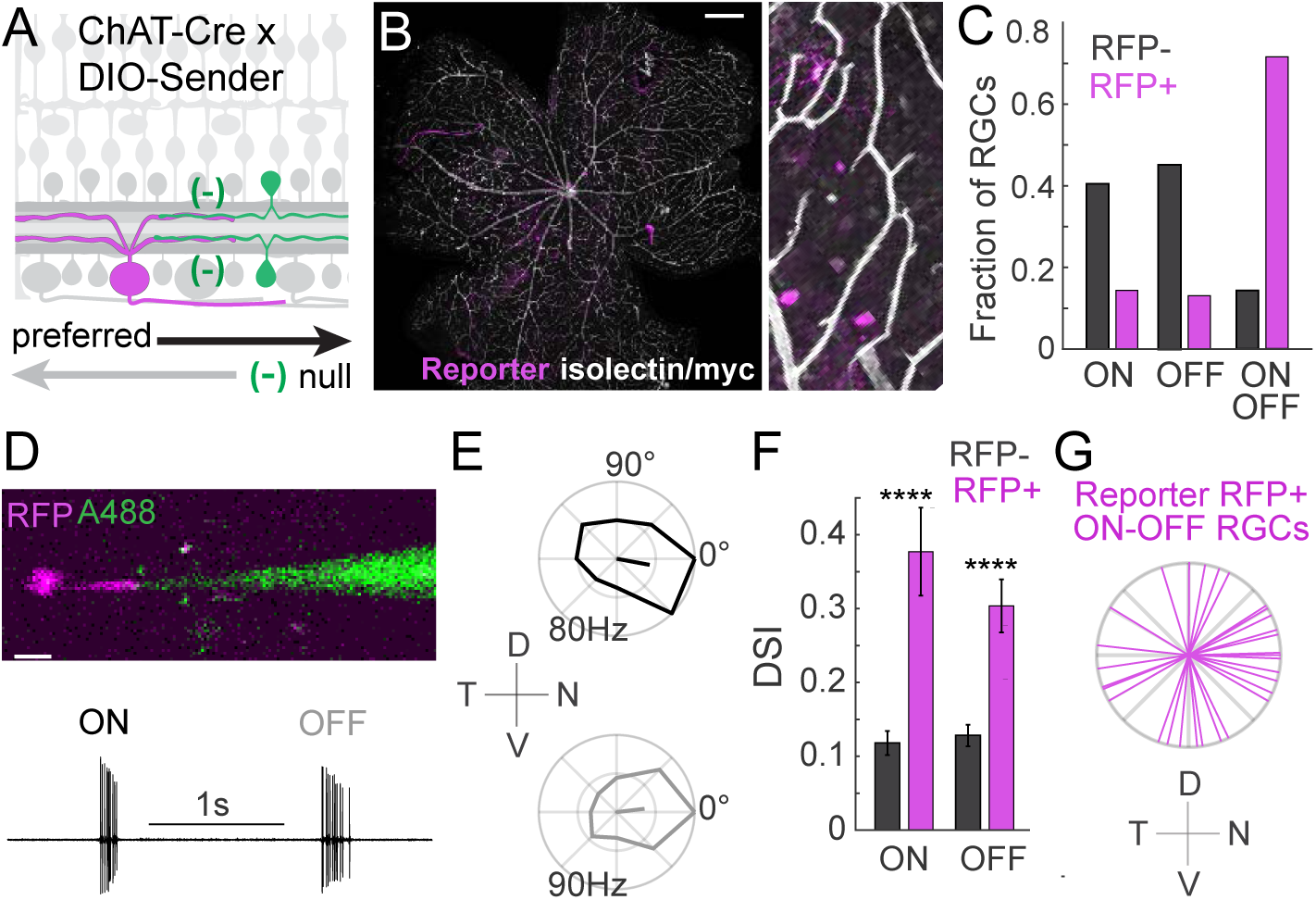
Functional recordings of TRACR-positive cells confirm direction-selective identity. **(A)** Schematic of TRACR strategy in *ChAT-Cre* retinas showing Sender-positive Starburst amacrine cells (green), Receiver+ ACs and RGCs (grey), and Reporter labelled DSGCs (RFP, magenta). Recorded RFP+ cells are predicted to show direction-selective responses to preferred direction (black). Mice received AAV intravitreal injections at 3-4 postnatal weeks and harvested at 2 months of age; AAV.TRE-mRuby and *Ai63* mice were used for Reporters. **(B)** Wholemount retina stained with antibodies against RFP, isolectin to label blood vessels, and anti-myc to mark Receiver-labelled retinal neurons. **Right:** High power image shows a typical field. **(C)** Fraction of Reporter RFP-positive (RFP+, n = 32 RGCs) and RFP-negative (RFP-, n= 42 RGCs) with ON, OFF, and ON-OFF responses to full field flash stimuli. **(D)** Two-photon image of alexa488 filled electrode (A488) and RFP+ RGC (top) and its spike responses evoked by a full field flash (bottom). **(E)** Polar plot of RFP+ RGC firing evoked by a bright bar moving in 8 directions. ON responses evoked by the leading edge of the bars; OFF responses evoked by the trailing edge. Normalized firing vector showing direction selective index (DSI) and angular preference also shown. **(F)** Average ON and OFF DSIs computed from recordings from RFP+ (n=32 RGCs) and RFP-cells (n=42 RGCs), like shown in E. Data show mean ± SEM. p < 0.05, t-test. **(G)** Polar plot showing angular preference from RFP+ ON-OFF RGCs. Data come from recordings in over 30 retinas from 25 mice. Scale bars: 1mm (B), 20 µm (D).

To examine this idea, we analyzed RGC responses to bright bars passed over their receptive field center moving in 8 different directions. As expected, many ON-OFF RFP-positive RGCs fired brief bursts of spikes when the bright bar’s leading edge entered the receptive field (ON) and again when the bar’s trailing edge exited the receptive field (OFF, Figure 3D). Polar plots of firing rate from these RFP-positive ON-OFF RGCs were often asymmetric indicating that these neurons fired most to bars movement in a particular direction (Figure 3E). Average ON- and OFF-direction selective indices computed from all RFP-positive and RFP-negative RGCs confirmed this picture and showed significantly higher average Direction Selectivity Index (DSI) for RFP-labelled neurons (Figure 3F). Finally, we plotted angular preferences aligned to the cardinal poles of retina for only ON-OFF RFP-positive RGCs and saw biased tuning towards dorsal, ventral, nasal and temporal directions (Figure 3G), consistent with prior reports of the directional tuning of ON-OFF DSGCs in the central part of the mouse retina (Vaney et al. 2012). These results support our anatomical experiments and show that TRACR outputs from SACs predominantly label ON-OFF direction selective RGCs.

Altogether, these data provide anatomical, molecular and functional evidence that TRACR labels the postsynaptic targets of genetically defined retinal neurons. They also highlight TRACR’s selectivity and utility for tracing local connections, including those of interneurons, where Senders and Receivers are expressed by physically proximate neurons.

### TRACR signals specifically across photoreceptor–bipolar cell synapses

To evaluate the sensitivity and specificity of TRACR signalling, we tested the system in the photoreceptor–ON bipolar cell circuit, a well-characterized synapse in which all contacts are confined to the outer plexiform layer (OPL), making them straightforward to visualize. We created a Sender-AAV with a photoreceptor-specific promoter (ProC1, (Jüttner et al. 2019)) to target NRX-GFP expression in rod and cone photoreceptors, and a Receiver-AAV with an ON bipolar-specific promoter to drive expression in ON cone and rod bipolar cells (Grm6[4x] promoter (Lagali et al. 2008)) (Figure 4A). To enable broad reporter expression in downstream cells, we used Ai63-TRE-tdTomato TIGRE reporter mice, which showed no detectable background expression in retina (Ai63-tdTomato; Daigle et al., 2018). TRACR AAVs were injected into the retinas of postnatal day (P)5 Ai63-tdTomato reporter mice. In initial tests, we observed RFP in Ai63 retinas injected with the hSyn-Receiver-AAV alone, suggesting ligand-independent activation of the Receiver 1.0 in Ai63 mice. We therefore replaced the Receiver with a modified synNotch-RAM7 to minimize background or leaky activation (Receiver 2.0) (Figure 1B). By restricting Sender and Receiver 2.0 expression to known synaptic partners, we evaluated whether TRACR reports postsynaptic partner identity and connectivity, rather than cellular proximity alone, and whether signaling is reversible and dependent on intact synapses.

**Figure 4.**
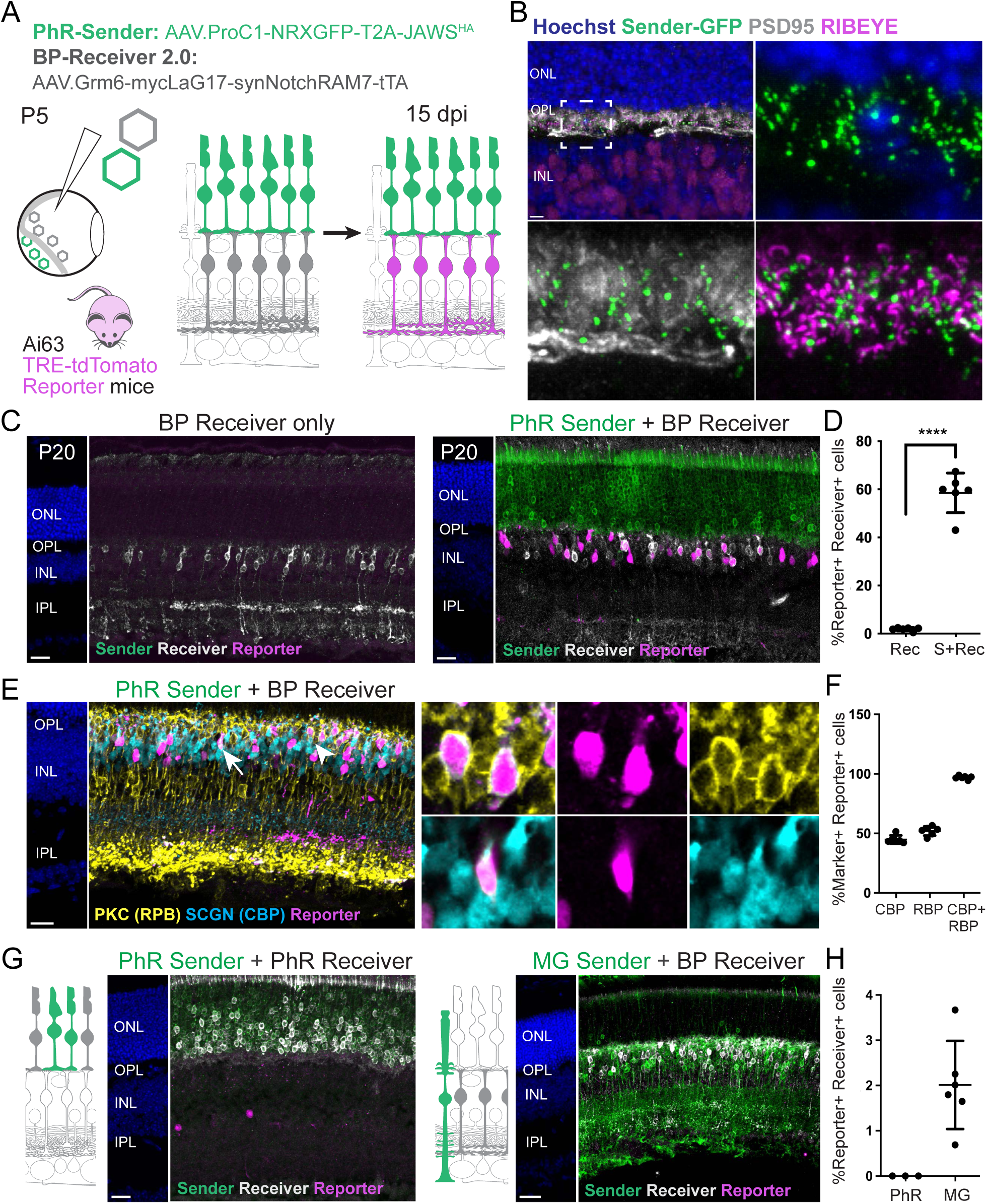
TRACR reports photoreceptor to bipolar cell connections. **(A)** Schematic of the TRACR strategy and predicted activation pattern at the photoreceptor–bipolar cell synapse. At P5, retinas of *Ai63* TRE-tdTomato Reporter mice were co-infected with a AAV-Sender driving NRX–GFP expression in photoreceptors (subretinal delivery of AAV.ProC1-NRXGFP-T2A-JawsHA), and AAV-Receiver 2.0 driving synNotch-RAM7 expression in ON-bipolar cells (intravitreal delivery of Receiver 2.0, AAV.Grm6-mycLaG17-synNotchRAM7-tTA). Synaptic contact between Sender- and Receiver 2.0-expressing cells induces RFP Reporter expression in Receiver-positive cells, analyzed at P20 (∼15 days post-infection, dpi). **(B)** Retinal cross-section showing localization of the NRX–GFP ligand in photoreceptor terminals (green, GFP fluorescence without amplification), in proximity to presynaptic markers Ribeye (magenta) and PSD95 (white). **(C)** TRACR activation requires Sender and Receiver 2.0 expression. Representative P20 retinal images show nuclear layers (Hoechst, blue; left panel), and stainings for Sender (GFP, green), Receiver (myc, white), and tdTomato Reporter (RFP, magenta). **Left:** Receiver 2.0-only controls show minimal ligand-independent reporter activation. **Right:** Co-injection of Sender and Receiver results in robust reporter activation confined to Receiver-expressing bipolar cells (middle). **(D)** Percentage of Reporter-positive Receiver-expressing cells in Receiver only (Rec) and Sender + Receiver (S + Rec) conditions. Each data point represents the mean of three sections from a single retina, 6 retinas per condition. Data are shown as mean ± SD. ****p < 0.0001, Welch’s t-test. **(E)** TRACR detects connections to rod and cone bipolar cells. Sender- and Receiver-infected retinas from P20 *Ai63* mice show RFP (magenta) labeling among rod bipolar cells (RBP; PKC, yellow; arrow), and cone bipolar cells (CBP; SCGN, cyan; arrowhead). Insets show examples of Reporter-positive RBPs and CBPs. **(F)** Percentage of marker-positive RFP-positive cells. Data are shown as mean ± SD. n= 6 retinas, 3 sections per retina. **(G)** Ligand proximity alone is insufficient to activate TRACR. **Left:** Schematic and representative retinal sections showing photoreceptor (PhR)-driven Sender (GFP, green; AAV.ProC1-NRXGFP) and Receiver expression (myc, white; AAV.ProC1-mycLaG17-synNotchRAM7-tTA). **Right:** Schematic and representative sections showing Müller glia-driven Sender (AAV.ProB2-NRXGFP) and bipolar-driven Receiver expression. **(H)** Percentage of Reporter-positive Receiver-expressing cells. Data are shown as mean ± SD. N = 3 retinas (PhR, Photoreceptor Sender/Receiver), 6 retinas (MG, Müller glia Sender), 3 sections per retina. Scale bars: 5 μm (B), 20 μm (C, E, G).

A key requirement for TRACR signalling is presynaptic localization of the NRX-GFP in Sender-expressing cells. Unamplified GFP fluorescence was concentrated at photoreceptor terminals in the OPL and colocalized with presynaptic proteins Ribeye and PSD95 (Figure 4B). In subsequent experiments, anti-GFP staining identified Sender-positive photoreceptors but showed a broader distribution within the cells, consistent with antibody detection of a low-level intracellular protein (Figure 4C). Together, these observations confirm ligand enrichment at presynaptic sites, satisfying a prerequisite for TRACR activation.

We next verified Grm6-driven Receiver 2.0 expression in bipolar cells. Immunostaining at P20 showed that the myc-tagged Receiver was confined to cells in the apical region of the INL displaying bipolar-like morphologies, consistent with Grm6 promoter activity in bipolar cells (Figure 4C). Approximately 83% of Receiver-positive cells were immunoreactive for Gαo, a pan–ON bipolar cell marker. The remaining ∼11% of Gαo-negative Receiver-positive cells expressed Chx10, a pan-bipolar cell marker (Supp. Figure 2). Thus, Receiver expression is largely restricted to ON bipolar cells, and nearly all Receiver-positive cells belong to the bipolar cell lineage. These findings demonstrate selective and efficient expression of TRACR components within the photoreceptor–bipolar cell circuit.

To determine whether TRACR activation depends on Sender-Receiver interaction, we compared reporter signal at P20 in Ai63-tdTomato mice injected with AAV Receiver alone versus those co-injected with AAV Sender and Receiver. Receiver-only injections produced minimal reporter signal in bipolar cells (1.9 % of Receiver-positive cells are RFP-positive), indicating low ligand-independent activation (Figure 4C,D). In contrast, co-injection of Sender and Receiver produced robust RFP reporter activation that was restricted to Receiver-expressing bipolar cells (58.5%; Figure 4C,D). These data demonstrate that TRACR activation is confined to Receiver-expressing cells and depends on the presynaptic ligand. Finally, Reporter-positive cells co-expressed either PKC, a marker of rod bipolar cells, or SCGN, a marker of cone bipolar cells (Figure 4E). Quantitative analysis confirmed significant TRACR activation in rod and cone bipolar populations, and that nearly all Reporter-positive cells were bipolar cells (Figure 4F). Therefore, TRACR detects synaptic connectivity across both rod and cone pathways.

To determine whether TRACR detects established synapses in the mature retina, we delivered the Sender AAV subretinally at P45 into eyes that had received the Receiver intravitreally at P5, as efficient bipolar-driven Receiver expression could not be achieved following adult delivery. After fifteen days, Reporter activation was again observed in Receiver-positive bipolar cells, and at levels comparable to those following neonatal virus delivery (Supp. Figure 2). Ligand-independent reporter activity in Receiver-only controls remained similarly low (Supp. Figure 2). Although Receiver expression was also observed in amacrine cells located in the basal INL at these timepoints, these cells did not activate the reporter, further confirming that direct Sender inputs are required for TRACR activation (Supp. Figure 2). Together, these experiments demonstrate that TRACR effectively labels postsynaptic partners in both developing and adult retinal circuits.

We tested similar Receiver 2.0; Ai63 TRACR combinations in the starburst amacrine cells in the retina, and in forebrain regions, where significant ligand-independent activation of Receiver 1.0 was also observed in Ai63 mice. In contrast to the inducibility of this system at the photoreceptor-bipolar synapse, Receiver 2.0 produced a few RFP-positive SAC targets in ChAT-Cre; Ai63 retinas in wholemount (Figure 3) or cryosections (Supp. Figure 3). We next tested TRACR with Receiver 2.0 in the brain, in a well-defined projection from the secondary auditory cortex to the lateral amygdala and striatum (LeDoux et al. 1991; Tsukano et al. 2019). Stereotaxic delivery of Sender-GFP AAVs to secondary auditory cortex induced Ai63-RFP reporter labeling in corresponding Receiver 2.0-expressing target regions, compared to sham-Sender AAV controls (Supp. Figure 4). Thus, the Receiver 2.0; Ai63 configuration increases specificity, although its sensitivity and inducibility appear to vary across synapse types.

Our next questions focused on validating that TRACR activation at the photoreceptor-bipolar cell synapse reflects transsynaptic interactions, rather than proximity between the Sender NRX-GFP ligand and the Receiver 2.0 receptor. We used two complementary strategies in which TRACR activation between adjacent cells, but not across synapses, would be expected to produce Reporter signal if proximity alone were sufficient. First, we expressed both Sender and Receiver in photoreceptors. We did not detect any reporter expression (Figure 4G,H). However, the co-expression of GFP ligand and synNotch receptors within the same cell could also cause *cis-*inhibition of TRACR activation, as reported for synNotch and endogenous Notch signaling (Morsut et al. 2016). Second, we expressed the Sender in Müller glia and the Receiver in bipolar cells. We generated a Müller glia–specific AAV-Sender (using ProB2 enhancer (Jüttner et al. 2019)) and confirmed its expression in Müller glia by co-immunostaining with Sox9 (Supp. Figure 5). Müller glial processes ensheath photoreceptor synapses (Burris et al. 2002; Williams et al. 2010), placing the Sender ligand in proximity to Receiver-expressing bipolar dendrites. Despite this anatomical proximity, reporter activation was limited to a few cells (Figure 4G,H). Together, these experiments demonstrate that TRACR is not activated by mere spatial proximity but instead requires presynaptic Senders in photoreceptors and postsynaptic Receivers in bipolar cells, establishing its specificity for transsynaptic contacts.

**Figure 5.**
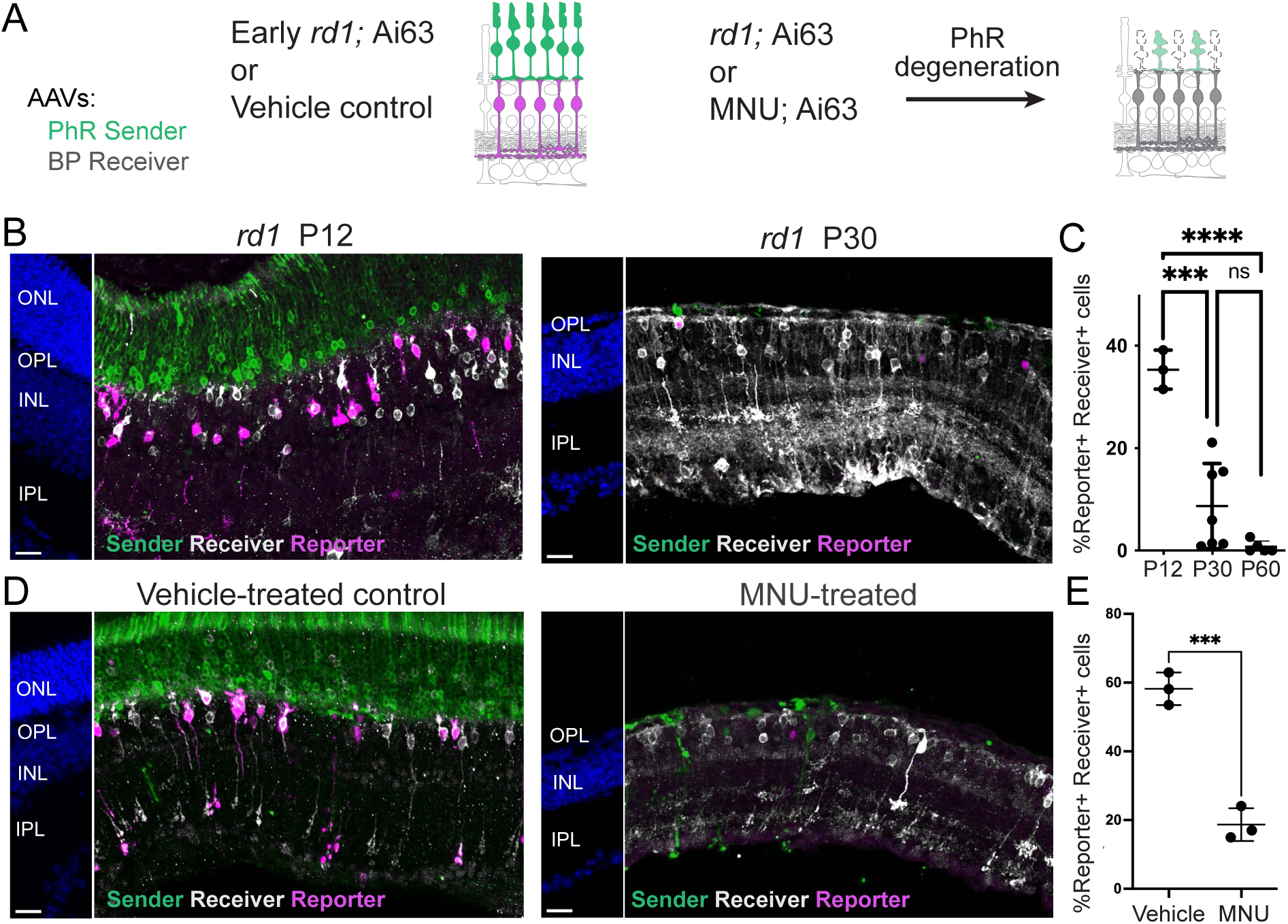
TRACR activation requires photoreceptor survival. **(A)** Predicted TRACR activation pattern following photoreceptor degeneration in *rd1;Ai63* and MNU-treated *Ai63* mice. Mice were infected at P5 with AAV.PhR-Sender and AAV.BP-Receiver (as in Figure 4A) and analyzed for reporter induction prior to and after photoreceptor degeneration. **(B)** TRACR activation is progressively lost during photoreceptor degeneration in *rd1* retina. Representative images from TRACR-infected *rd1;Ai63* retina showing Sender (GFP, green), Receiver (myc, white), Reporter (TdTomato, magenta), and nuclei (Hoechst, blue). P12 and P30 correspond to early and late-stage rod degeneration, visible by reduced ONL (blue panels). **(C)** Percentage of Reporter-positive Receiver-expressing cells at the indicated time points. Each data point represents the mean of three sections from a single retina. n = 3 retinas (P12), 7 retinas (P30), 5 retinas (P60). Data are shown as mean ± SD. ****p < 0.0001, *** p = 0.0001, one-way ANOVA, with Tukey pairwise comparisons. **(D)** TRACR activation is reduced in an acute photoreceptor degeneration model. Representative sections from TRACR-infected *Ai63* retinas and treated with MNU or vehicle at P20, then harvested at P38. MNU-treatment leads to reduced ONL due to PhR loss (Hoechst, blue). Robust Reporter activation (tdTomato, magenta) in Receiver-labelled BPs (myc, white) is observed in vehicle-treated controls but reduced in MNU-treated retinas. **(E)** Percentage of Reporter-positive Receiver-expressing cells in control and MNU-treated mice. N = 3 sections per retina, 3 retinas per condition. Data are shown as mean ± SD. p =0.0013, Welch’s t-test. Scale bars: 20 μm (B, D)

### TRACR reports loss of photoreceptor–bipolar cell connectivity following photoreceptor degeneration

Most tracing tools rely on recombinase-driven reporters that permanently mark synaptic partners, so that labels persist even after synapses are eliminated or remodeled. TRACR instead uses the reversible tTA–Tet-Off system, which should require ongoing synNotch cleavage and transcription driven by intact synaptic contact. We therefore asked whether TRACR activation is lost upon loss of photoreceptor–bipolar cell connectivity, a fundamental requirement for using this system to evaluate disease models or genetic perturbations that disrupt synapses.

We tested TRACR activation in two photoreceptor degeneration models. We first used the rapid genetic rod degeneration model, *rd1* (Bowes et al. 1990), and delivered the photoreceptor-specific Sender and bipolar-specific Receiver AAVs in retinas of P5 *rd1;* Ai63 animals (Figure 5A). At P12, prior to substantial photoreceptor loss, Sender and Receiver expression as well as TRACR Reporter activation were robust, indicating that compromised photoreceptors in this model can still express the constructs and induce TRACR signaling across synaptic contacts (Figure 5B,C). By P30, when most rods have degenerated, Reporter expression was significantly reduced, and by 2 months it was nearly undetectable (Figure 5B,C). Because *rd1* is an early-onset degeneration model, we also assessed acute photoreceptor loss in adult retinas. We induced degeneration with MNU, which rapidly eliminates photoreceptors (Petrin et al. 2003). In MNU-treated eyes of adult mice, Sender-positive cells were lost and TRACR activation was significantly reduced compared to vehicle-treated controls (Figure 5D,E). Thus, TRACR signal declines as synaptic contacts are eliminated, and the readout depends on the continued presence of presynaptic photoreceptors. Together, these results show that TRACR reversibility enables analysis of synapse disruption in disease and injury models.

### TRACR detects synaptic disruption in genetic and activity-dependent models

We then investigated whether TRACR can detect synaptic disruption, as this would enhance its utility for assessing how disease models or genetic modifications impact connectivity. We evaluated TRACR activation in *Nrl-Cre; Elfn1^fl/fl^*;Ai63 mice with a rod photoreceptor-specific inactivation of *Elfn1* (*Rod-Elfn1* cKO). ELFN1 is a rod-specific transsynaptic adhesion molecule essential for the establishment of rod-to-rod bipolar cell synapses (Cao et al. 2015) (Figure 6A). In this instance, reporter induction was significantly reduced in Rod-*Elfn1* cKO mice compared to controls, despite the persistence of both cell types (Figure 6B,C). To determine whether loss of TRACR activation was specific to *Elfn1* disruption or reflected a broader sensitivity to synaptic failure, we also examined *Nrl-Cre; R26-LSL-TeNT*; Ai63 mice in which tetanus toxin is developmentally expressed in rods (*Rod-TeNT*) (Zhang et al. 2008) (Supp. Figure 6). This intervention abolishes SNARE-mediated synaptic vesicle release in rods, and disrupts synapse formation between rods and ON rod bipolar cells (Cao et al. 2015). Reporter induction in rod bipolar cells was significantly decreased in Rod-TeNT mice compared to controls (Supp. Figure 6). Interestingly, most of the remaining RFP-positive cells were SCGN-positive cone bipolar cells in both the *Rod-Elfn1* cKO and *Rod-TeNT* mice, consistent with preserved cone-to–cone bipolar connectivity when rod pathways are selectively impaired (Figure 6D,E, Supp. Figure 6). Together, these findings show that in two independent models of rod synapse dysfunction, TRACR selectively reports intact cone connectivity.

**Figure 6.**
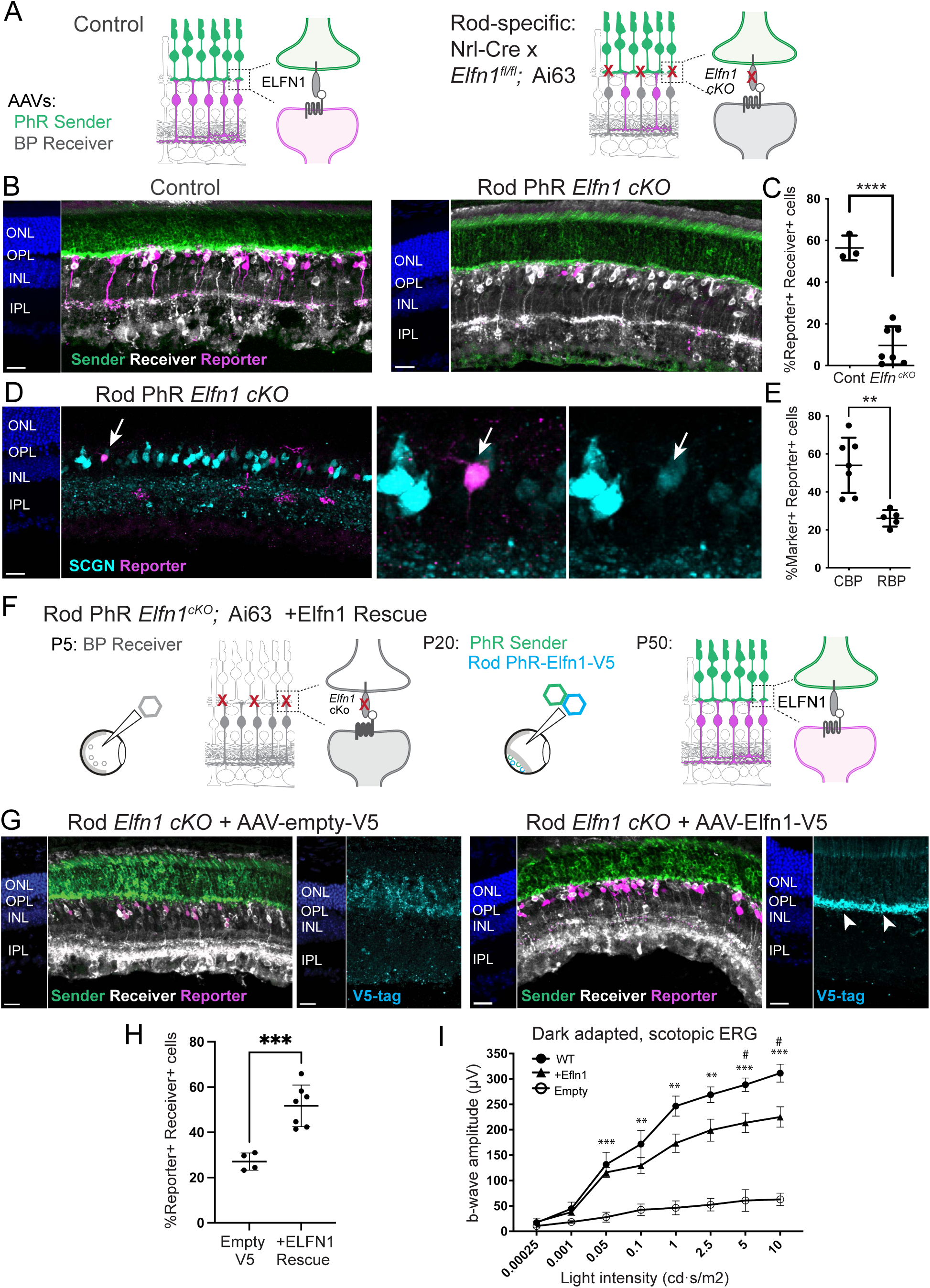
TRACR activation requires intact photoreceptor-to-bipolar cell synapses. **(A)** Strategy to disrupt transsynaptic connectivity via rod-specific deletion of *Elfn1* (*Nrl-Cre;Elfn1flox; Ai63*). Mice were infected with AAV.PhR-Sender and AAV.BP-Receiver at P5 and analyzed at P20. TRACR induction is predicted to be maintained for PhR to cone bipolar cell synapses (magenta) but abolished between rod PhR to RBP connections (grey). **(B)** TRACR activation is reduced when rod–bipolar synapse formation is disrupted. Representative retinal sections from P20 control (*Elfn1fl/fl;Ai63*) and *Nrl-Cre; Elfn1fl/fl;Ai63* mice immunostained for Sender (HA, green), Receiver (myc, white), Reporter (TdTomato, magenta), and nuclei (Hoechst, blue). **(C)** Percentage of Reporter-positive Receiver-expressing cells. n= 3 sections per retina, 3 retinas (control) and 7 retinas (*Nrl-Cre; Elfn1fl/fl; Ai63*). Data are shown as mean ± SD. ****p < 0.0001. **(D)** TRACR activation is preserved in cone bipolar cells (CBP) when rod–bipolar connectivity is disrupted. Representative retinal sections from *Nrl-Cre; Elfn1fl/fl; Ai63* mice show Reporter expression (tdTomato, magenta) within CBPs (SCGN, cyan; arrows). **(E)** Percentage of CBP (SCGN-positive) or RBP (PKC-positive) Reporter-positive cells. n=3 sections per retina, 7 retinas per SCGN marker, 5 per PKC marker. Data are shown as mean ± SD. **p = 0.002. **(F)** Strategy to restore rod-bipolar synaptic connectivity in *Nrl-Cre; Elfn1fl/fl; Ai63* mice and predicted TRACR activation pattern. Retinas received AAV.BP-Receiver at P5, and co-injected subretinally with AAV.PhR-Sender and either rod-specific AAV.ProA6-Empty-V5 (control) or AAV.ProA6-Elfn1-V5 (+ELFN1). **(G)** TRACR activation is restored following ELFN1 re-expression. Representative retinal sections from *Nrl-Cre; Elfn1fl/fl; Ai63* mice infected with AAV-empty-V5 (left) or AAV. ProA6-Elfn1-V5 (right), showing Sender (HA, green), Receiver (myc, white), Reporter (tdTomato, magenta), V5 (cyan) and nuclei (Hoechst, blue). Arrowheads confirm ELFN1-V5 localization within rod terminals in the OPL. **(H)** Percentage of Reporter-positive Receiver-expressing cells. n= 3 sections per retina, 4 control (*Nrl-Cre; Elfn1fl/fl; Ai63* + AAV-Empty-V5) and 7 rescue (Nrl-Cre; Elfn1fl/fl; Ai63 + AAV-Elfn1-V5). Data are shown as mean ± SD. ***p < 0.0002. **(I)** Scotopic ERG b-wave amplitudes plotted as a function of flash intensity in wild-type, control and +ELFN1 rescue eyes. Data are shown as mean ± SEM. n = 5 wild-type, 4 control, and 7 +ELFN1 eyes. Statistical significance was determined by a two-way ANOVA followed by Tukey’s multiple comparisons test. Asterisks indicate significant differences between the +ELFN1 and control groups (**p <0.01, ***p <0.001); hashtags indicate significant differences between the WT and Rescue groups (#p <0.05). Scale bars: 20 μm (B,D,G).

### TRACR detects restoration of synaptic connectivity

A key application of transsynaptic tracing tools is the ability to monitor circuit repair following synapse disruption. Previous studies showed that re-expression of ELFN1 restores rod–rod bipolar cell synapses and recovers rod-mediated visual responses in *Elfn1* mutant mice (Cao et al. 2022), providing an opportunity to test whether TRACR can detect synapse restoration *in vivo*. To address this, Rod-*Elfn1 cKO* mice were infected with the ON Bipolar-Receiver AAV at P5, followed by subretinal co-delivery at P20 of PhR-Sender together with a rod-specific ProA6 AAV expressing V5-tagged ELFN1, or a control ProA6 AAV expressing V5 alone (Figure 6F). ELFN1 re-expression was confirmed by immunostaining for the V5 epitope (Figure 6G; Supp. Figure 6). Thirty days after AAV delivery, ELFN1-expressing retinas exhibited significantly increased Reporter activation compared with retinas injected with a control vector (Figure 6G, H). Consistent with restoration of rod bipolar circuit function (Cao et al. 2022), ELFN1-treated eyes also displayed recovery of scotopic ERG responses (Figure 6I). Together, these findings demonstrate that TRACR can detect restoration of a defined synaptic connection following genetic rescue and identify the postsynaptic cell types engaged during circuit repair.

In summary, TRACR detects functional rod-to–rod bipolar cell synapses and fails to activate when synapses are lost, silent, or absent, demonstrating that the system provides a reversible readout of *bona fide* synapses and reports genuine alterations in photoreceptor–bipolar connectivity. These experiments further highlight the potential for TRACR to label synaptic partnerships during circuit remodeling and repair across sensory, local and projection pathways in the mouse nervous system.

## DISCUSSION

A persistent limitation in circuit mapping is the lack of broadly adoptable anterograde tracing tools that provide separable genetic access to both sides of a connection. We developed TRACR, an AAV-deliverable toolkit that adapts synthetic Notch signaling to convert presynaptic ligand binding into tTA-TRE driven reporter expression in postsynaptic cells. Across multiple synapses in the mouse visual system, TRACR labeled targets downstream of sensory neurons, inhibitory interneurons, and long-range projection neurons, and distinguished partners within local circuits. Because TRACR signaling requires intact synaptic organization, its transcriptional output provides a temporally sensitive readout of connection status, including changes following photoreceptor degeneration in *rd1* mice and ELFN1-mediated rescue of rod synapse assembly. TRACR components are modular and sufficiently compact for standard AAV vectors, and enable transneuronal control of gene expression that can be integrated with existing genetic markers, sensors, or optogenetic actuators. Together, these results establish TRACR as a broadly deployable platform for anterograde transneuronal tracing and longitudinal circuit analysis in mammals.

TRACR represents the first mammalian implementation of a ligand–receptor tracing strategy that couples synaptic contact to transcription. Conceptually, it parallels TRACT and Trans-Tango in *Drosophila*, which similarly convert synaptic contact into gene expression using synNotch or engineered GPCR signalling, respectively (Huang et al. 2017; Talay et al. 2017). A key distinction is that synNotch systems rely on endogenous proteases such as γ-secretase, whereas Trans-Tango uses reconstituted TEV protease recruited to an engineered receptor. In zebrafish, Trans-Tango has been combined with optogenetic and calcium imaging tools to probe functional coupling *in vivo* (Coomer et al. 2025). TRACR was also designed to support all-optical experiments. In the current Sender configuration, however, ChR2 after a 2A cleavage peptide did not reliably evoke strong photocurrents in SACs. Alternatively, the Sender AAV containing a Flpe recombinase could be delivered along with Flp-dependent opsins. Studies are underway to effectively integrate optogenetic activation.

### TRACR requires an intact synaptic structure

Benchmarking TRACR at the photoreceptor–bipolar synapse showed that induction is Sender-dependent, with minimal ligand-independent activation, and synapse-dependent, as signaling is lost when input synapses are disrupted. Altough these experiments do not define the precise nanoscale site of Sender–Receiver interaction, several observations suggest that signaling occurs within specialized synaptic domains rather than through proximity alone. The Sender was strongly enriched in photoreceptor and RGC presynaptic terminals, consistent with the ability of neurexin-beta intracellular domains to promote surface delivery and presynaptic localization (Fairless et al. 2008; Gokce and Südhof 2013). Photoreceptor synapses comprise an extended transsynaptic membrane interface containing ELFN1 trans-synaptic organizers, mGluR6-associated signaling machinery, and dystrophin-glycoprotein complexes that are distributed along the invaginating rod bipolar dendrites rather than confined to the ribbon active zone (Tsukamoto and Omi 2022). Sender–Receiver interactions may occur within this broader specialized synaptic domain rather than at a position directly opposite the ribbon. Consistent with this interpretation, TRACR signaling was abolished in rod-specific *Elfn1* conditional mutants and in tetanus toxin-expressing rods (*Rod-TeNT*). In both models, rod terminals target the OPL and retain aspects of presynaptic specializations but postsynaptic differentiation of rod ON-bipolar targets, including dendritic invagination, is disrupted (Cao et al. 2015). Because blocking vesicular fusion in *Rod-TeNT* mice impairs ELFN1–mGluR6 signaling and synaptic maturation (Cao et al. 2015), these experiments do not distinguish whether loss of TRACR signaling reflects impaired synaptogenesis, reduced synaptic transmission, or both.

We posit that presynaptic Sender enrichment raises the ligand concentration above the threshold for synNotch activation and reporter induction in connected partners, but not in neighbouring processes. Supporting this model, Sender-expressing Müller glia did not drive reporter induction in adjacent Receiver-expressing bipolar cells. Importantly, TRACR activation was lost following photoreceptor degeneration and *Elfn1* deletion, and restored following ELFN1-mediated synapse rescue. Together, these findings indicate that TRACR reports organized synaptic connectivity rather than spatial proximity alone.

### Tracing advances offered by TRACR

TRACR is distinct among mouse transneuronal tracers as it uses a synthetic ligand–receptor to report a synaptic interaction into a postsynaptic transcriptional readout, rather than the transfer of viral particles or proteins. TRACR’s anterograde directionality is determined by the presynaptic targeting of the GFP ligand, whereas labeling is restricted to Receiver-expressing downstream cells. This contrasts with anterograde viral and WGA-based approaches, whose transfer mechanisms are less defined with potential for bidirectional or polysynaptic spread, and whose efficiency varies with cell type and circuit context (Zingg et al. 2017; Beier 2019; Xu et al. 2020; Zingg et al. 2020; Li et al. 2021; Tsai et al. 2022; Xiong et al. 2022). ATLAS provides an alternative strategy in which engineered proteins are released from presynaptic vesicles and endocytosed by postsynaptic cells through binding to neurotransmitter receptors, but it is currently limited to excitatory synapses (Rivera et al. 2025). By contrast, TRACR signals across both excitatory and inhibitory synapses, enabling mapping across diverse circuit organizations. Its TRE-driven output is also reversible, unlike the permanent recombinase-based labeling used by most anterograde approaches, creating opportunities to track changes in connectivity over time.

Unlike many viral approaches prone to cellular toxicity, TRACR was non-toxic and did not measurably perturb synapse integrity or function. We showed this in three ways. First, Sender and Receiver expression in multiple retinal cell types did not cause overt cell loss or retinal thinning. Second, TRACR signaling persisted in wild-type retinas over many weeks, whereas signaling declined or was absent following photoreceptor degeneration (*rd1*, MNU treatment), neurotransmission blockade (TeNT), or impaired synapse formation (*Elfn1 cKO*). Third, TRACR-positive retinal ganglion cells downstream of Sender-expressing SACs retained normal ON–OFF responses and direction tuning to moving stimuli. Together with its reliance on standard AAV delivery, these features make TRACR broadly accessible compared to pathogenic tracers that have greater biosafety and regulatory constraints.

An additional strength is TRACR’s modularity and genetic access to either side of the synapse. Sender expression can be restricted to defined presynaptic populations using Cre lines and the expanding repertoire of cell-type selective AAV enhancers and promoters (Hrvatin et al. 2019; Jüttner et al. 2019; Graybuck et al. 2021; Ben-Simon et al. 2025; Furlanis et al. 2025). This differs to approaches such as AAV1-Cre or YFV-17D that transfer Cre itself and limit presynaptic specificity (Zingg et al. 2017; Zingg et al. 2020; Li et al. 2021). For unbiased partner discovery, the Receiver and Reporter can be expressed broadly so that postsynaptic labeling is agnostic to target identity, provided that AAV infection is uniform. TRACR can also leverage tTA-dependent AAVs and transgenic mice to mark post-synaptic populations with reporters or genetically encoded indicators or effectors of choice (Daigle et al. 2018). The TRACR design is interchangeable with common Cre, Flp and other transcriptional systems, enabling highly customisable outputs, with the main practical constraint being AAV packaging capacity.

Finally, TRACR appears effective across neuron classes and synapse types, and it resolves local circuits, which remains difficult for most transsynaptic tools. In the visual system, TRACR labeled predicted postsynaptic partners of sensory neurons (photoreceptors) and GABAergic interneurons (amacrine cells). The retina’s inner plexiform layer (IPL) is a particularly stringent setting for testing TRACR in local wiring because processes from over 100 bipolar, amacrine, and ganglion cell types intermingle yet form stereotyped microcircuits. Within this dense neuropil, TRACR distinguished targets of starburst amacrine cells from those of broader amacrine populations. Proximity and co-stratification of processes are insufficient to infer connectivity in the IPL (Helmstaedter et al. 2013; Bae et al. 2018; Prigge et al. 2023), with clear examples of synapse partnering choices determined by recognition molecules and required for circuit output (Krishnaswamy et al. 2015; Rochon et al. 2021; Friedrichsen et al. 2024; Olguin et al. 2025). As the connectivity of many circuits remain unresolved in the retina and beyond, TRACR stands to be a valuable tool for mapping synaptic partnerships.

### Specificity, sensitivity, and limitations of TRACR

As with other AAV-based tools, TRACR performance is influenced by viral dose, infection efficiency, and expression levels. Consistent with prior reports of Tet-OFF leakiness and ligand-independent synNotch activation, we observed low-level reporter expression with Reporter AAV alone, and with Receiver (1.0) plus Reporter AAVs in the absence of Sender. Empirical titration of Receiver and Reporter AAVs minimized this background. Thus, appropriate controls and titration of each component should be conducted within each circuit of interest when using the AAV complement of TRACR.

Combining TRACR with Ai63-TITL-tdTomato reporter mice simplified the experimental design and did not produce tTA-independent reporter expression. However, the original synNotch (Receiver 1.0) produced ligand-independent activation in the retina and brain of Ai63 mice. Replacing Receiver with synNotch-RAM7 (Receiver 2.0), which contains a hydrophobic sequence native to the Notch core that suppresses ligand-independent activation (Yang et al. 2020), minimized reporter background in Ai63 controls. At the photoreceptor–bipolar synapse, Receiver 2.0 achieved high sensitivity, inducing Reporters in ∼60% of Receiver-positive cells in wildtype retinas, with variability likely reflecting infection efficiency. By contrast, the senstivity of Receiver 2.0 was severely diminished for tracing SAC targets in the inner retina, indicating that the inducibility of this TRACR combination varies with synapse type. The threshold for Receiver 2.0 activation may be determined by synaptic anatomy, Sender abundance, or the number of convergent Sender inputs. Increasing Sender expression with strong promoters, higher Sender-to-Receiver ratios (Malaguti et al. 2022), and longer infection periods can improve induction. A future direction will be to generate dedicated TRACR mouse lines with optimized Receivers that reduce expression variability and provide more consistent sensitivity and specificity across synapses and circuits.

Another limitation is that the GFP ligand uses neurexin-3beta intracellular sequences that promote surface delivery at axonal membranes and presynaptic enrichment but are not sufficient to exclusively restrict the ligand to synaptic interfaces (Fairless et al. 2008; Gokce and Südhof 2013; Klatt et al. 2021). Stable neurexin complexes at synapses is thought to require extracellular interactions with neuroligins through the neurexin LNS domain, but this domain risks ectopic synaptogenesis when overexpressed (Kim et al. 2011; Tsetsenis et al. 2014). Further strategies to improve the pre- and postsynaptic restriction of TRACR components are being explored. As with any transsynaptic method, TRACR outputs should be interpreted with the possibility of false positives and false negatives, and further validated using complementary anatomical and functional assays.

In summary, TRACR offers a unique, reversible transneuronal tracing method for unbiased, high-throughput identification of postsynaptic partners and longitudinal analysis of mammalian circuit connectivity. By coupling synaptic contact to transcription, TRACR can reveal connectivity gains and losses during development, disease, and genetic perturbations that alter synapse formation or maintenance.

## MATERIALS and METHODS

### Animal care and mouse lines

Animal studies were carried out in accordance with the guidelines of the Canadian Council on Animal Care, and under protocols approved by the Animal Care Committees of the Centre for Phenogenomics (TCP), Laboratory Animal Services (LAS) at the Hospital for Sick Children, University Health Network Research Institute (Toronto Canada) and the Goodman Cancer Research Facility Vivarium at McGill University (Montreal, Canada). All facilities are certified by the Canadian Council on Animal Care and while TCP, LAS, and University Health Network facilities registered under the Animals for Research Act of Ontario.

Mice were maintained on a C57/B6J background. Both male and female mice were used at all ages. The following mouse lines were purchased from the Jackson Laboratory: ChAT-ires-Cre (B6;129S6-*Chat^tm2(cre)Lowl^/*J, Strain #:006410; RRID:IMSR_JAX:006410) (Rossi et al. 2011); Gad2-ires-Cre (*Gad2^tm2(cre)Zjh^*/J; Strain #:010802; RRID:IMSR_JAX:010802) (Taniguchi et al. 2011); Vglut2-ires-Cre (B6J.129S6(FVB)-*Slc17a6^tm2(cre)Lowl^*/MwarJ, Strain #:028863; RRID:IMSR_JAX:028863) (Vong et al. 2011); Nrl-Cre (C57BL/6J-Tg*(Nrl-cre)1Smgc*/J, Strain #:028941; RRID:IMSR_JAX:028941) (Brightman et al. 2016); and *Rd1* mice with retinal degeneration phenotype (C57BL/6J-*Pde6b^rd1-2J^*/J (Strain #:004766; RRID:IMSR_JAX:004766).

Ai63(TIT-tdTomato) are tTA-dependent reporter mice expressing tdTomato from the TRE-tight promoter, and were created by targeted insertion at the Igs7 TIGRE locus (B6.Cg-*Igs7^tm62.2(tetO-tdTomato)Hze^*/J)(Daigle et al. 2018). Ai63 mice were obtained from the Allen Institute for Brain Science (a kind gift from Dr. Hongkui Zeng, Seattle, USA). R26LSL-TeNT (Gt(ROSA)26Sor*^tm2(GFP/tetX)^*Gld) (Zhang et al. 2008) animals were a kind gift from Dr. Michel Cayouette (Institut de Recherches Cliniques de Montréal, Montreal, Canada). The *Elfn1^fl/fl^* mice were described previously (Cao et al. 2015).

### Plasmid cloning

Sender plasmids were generated as follows: To create the NRX-GFP ligand, codon-optimised sequences of the transmembrane and cytoplasmic domains of the human Neurexin-3β (NP_620426.2) followed by the membrane trafficking signal of human Kir2.1 (NP_000882.1) (Gradinaru et al. 2010) were synthesized as gBlocks (Integrated DNA Technologies). eGFP was amplified from pCAGIG (Matsuda and Cepko 2004) with the addition of the N-terminal signal peptide sequence of rat β2 acetylcholine receptor (Zhao et al. 2008). To create pAAV.hSyn-DIO-Nrx3bGFP-T2A-smFP^HA^-WPRE-pA, fragments were fused using Gibson assembly (# E2611S, New England Biolabs) into a pCAG-driven backbone to create a transient vector. Nrx3bGFP and smFP^HA^ derived from pCAG-smFP_HA (gift from Loren Looger; Addgene plasmid # 59759; http://n2t.net/addgene:59759; RRID:Addgene_59759) (Viswanathan et al. 2015) were PCR amplified with overlapping tails to insert a T2A sequence, and assembled between the loxP sites of a linearized hSyn-containing AAV backbone by Gibson assembly (pAAV.hSyn-DIO-hM3D(Gq)-mCherry, a gift from Bryan Roth, Addgene plasmid # 44361; http://n2t.net/addgene:44361; RRID:Addgene_44361) (Krashes et al. 2011). To create a Sender with a channelrhodopsin marker (pAAV.hSyn-DIO-Nrx3bGFP-T2A-hChR2(H134R)-eYFP-WPRE-pA), the smFP^HA^ insert was replaced with hChR2(H134R)-eYFP (Addgene plasmid #26973, a gift from Karl Deisseroth; http://n2t.net/addgene:26973; RRID:Addgene_26973). To create CBh-driven Senders (pAAV.CBh-DIO-Nrx3bGFP-T2A-ChR2(H132R)-eYFP-WPRE3-pA), the Sender insert was subcloned into AAV backbone (AiP11839-pAAV.hSyn1-SYFP2-10aa-H2B-WPRE3-BGHpA, Addgene plasmid # 163509; http://n2t.net/addgene:163509; RRID:Addgene_163509a; gift from Allen Institute for Brain Science & Jonathan Ting) (Graybuck et al. 2021) and the hSyn promoter was replaced with CBh (from pAAV.CBh-DIO-LifeAct-eGFP-WPRE) (Ing-Esteves and Lefebvre 2024). Sender plasmid with T2A-Flpo (pAAV.hSyn-DIO-Nrx3bGFP-T2A-Flpo-WPRE) allows expression of Flp-dependent vectors to complement Sender expression. pAAV.hSyn-DIO-Nrx3bGFP-T2A-Flpo-WPRE was created by amplifying a FlpO sequence from Addgene plasmid #26745 (gift from Rolf Zeller; http://n2t.net/addgene:26745; RRID:Addgene_26745) (Osterwalder et al. 2010) which was then inserted into pAAV.DIO-Sender-GFP backbone by Gibson Assembly.

To create retina cell-type-specific Senders, synthetic promoters developed by Jüttner *et al*. (Jüttner et al. 2019) were PCR amplified or subcloned from the following plasmids: pan-photoreceptor ProC1 (pAAV.ProC1-CatCh-GFP-WPRE, gift from Botond Roska, Addgene plasmid # 12593; http://n2t.net/addgene:125937; RRID:Addgene_125937); Müller Glia ProB2 (pAAV.ProB2-CatCh-GFP-WPRE, Addgene plasmid # 125922; http://n2t.net/addgene:125922; RRID:Addgene_125922). Amplified fragments were inserted into a linearized Sender-AAV backbone using standard restriction digest and ligation. The ChR2 marker was replaced by Jaws-KGC-ERT2 optogenetic channel (Chuong et al. 2014) tagged with hemagglutinin (HA). The Jaws-HA insert was synthesized (Bio Basic Inc., Markham, Canada) and inserted into the linearized backbone to create a series of Senders, pAAV.Pro(C1, B2 or A6)-Nrx3bGFP-T2A-Jaws-HA-WPRE3-pA.

The Receiver construct (pAAV.hSyn-LaG17-synNotch-tTA-WPRE-pA) was created by PCR amplification of sequences encoding the LaG17 nanobody, synNotch and tTA (Tet-OFF) from pHR-SFFV-LaG17-synNotch-TetR-VP64 (gift from Wendell Lim; Addgene plasmid # 79128; http://n2t.net/addgene:79128; RRID:Addgene_79128) (Morsut et al. 2016) along with an N-terminal myc epitope tag, and inserted into a linearized AAV.hSyn backbone. To reduce the ligand-independent activation observed in Ai63 TRE-tdTomato mice injected with AAV.Receiver only, we exchanged a synNotchRAM7 containing a hydrophobic sequence present in the endogenous Notch (QHGQLWF) (Yang et al. 2020). SynNotchRAM7 was synthesized commercially (Biobasic, Canada) and inserted into the pAAV.hSyn-LaG17-tTA vector. To create the pAAV.Grm6[4x]-LaG17-synNotch RAM7-tTA for selective expression in rod bipolar and ON cone bipolar cells, we inserted the promoter sequence containing four tandem repeats of a 200 bp element of the metabotropic glutamate receptor 6 gene (pAAV.Grm6S[4X]-tdTomato plasmid was a gift from Dr. Daniel Kerschensteiner) (Lagali et al. 2008; Tien et al. 2017).

To create the pAAV.ProA6-Elfn1-V5-WPRE3-SL vector, an Elfn1 cDNA sequence (Cao et al. 2020) was codon-optimized and synthesized with a V5 epitope tag (GeneArt, ThermoFisher), and inserted into an pAAV.WPRE3-SL vector that includes the rod-specific ProA6 promoter (gift from Botond Roska, Addgene plasmid # 125890; http://n2t.net/addgene:125890; RRID:Addgene_125890). The empty vector control was constructed by fusing a CD8 signal sequence to V5, pAAV.ProA6-CD8-V5-WPRE3-SL.

### AAV administration

Recombinant adeno-associated virus (rAAVs) for the TRACR system used in this study were produced as AAV2/9 or AAV2.7M8 viral particles (∼1-5 x 10^12-13^ viral genome (vg)/mL) by Vigene BioSciences/Charles River (Rockville, MD, USA) or the Canadian Optogenetics and Vectorology Foundry (COVF) Viral Vector Core (Laval, Canada; RRID:SCR_016477). Viruses used and experiments are listed in Table S1.

The following plasmids were obtained from Addgene (Watertown, MA, USA), and AAV2/9 vectors were produced by the COVF Viral Vector Core: pAAV.hSyn-DIO-tdTomato-2A-9xHA-Synaptophysin (a gift from Bernardo Sabatini, Addgene plasmid # 163686; http://n2t.net/addgene:163686; RRID:Addgene_163686) (Chantranupong et al. 2020); pAAV.TRE-mRuby2 (a gift from Viviana Gradinaru, Addgene plasmid # 99114; http://n2t.net/addgene:99114; RRID:Addgene_99114) (Chan et al. 2017).

Intraocular injections of AAVs: AAVs were delivered into eyes of postnatal (P0-P4) or juvenile (P25 or older) mice by intraocular injection procedures described previously (Ing-Esteves et al. 2018; Gurdita et al. 2023). Postnatal pups were anesthetized by hypothermia by placing animals on ice until they were unresponsive to touch). Juvenile mice were anesthetised with 4% isoflurane and maintained on 2-2.5% isoflurane or with ketamine at 100 mg/kg of body weight for P20 and older. Animals were placed on a warming pad to maintain body temperature and fully recover from anesthesia before being returned to their cages.

For Sender and Receiver infections in the inner retina, intravitreal AAV delivery of juvenile mice, a 30-1/2 gauge needle (Becton Dickinson Mississauga, Canada) used to make a small hole just above the ora serrata, and 0.7-1.0 μl of rAAV mixed with Fast Green dye for visualisation was injected with a Hamilton syringe (7632-01) and 33 gauge blunt-ended needle (7803-05, Hamilton) into the intravitreal space. For Sender- and Receiver-AAV infection of photoreceptor or Müller glial cells in the outer retina, a subretinal injection was performed. Briefly, a 30G needle (BD, Mississauga, Canada) was used to make a small incision in the sclera at the nasal limbus, and the refluxed vitreous fluid was removed using Surgical spears (30-049, DeRoyal, Tennessee, US). A Hamilton syringe (80016, Chromatographic Specialties Inc., Brockville, Canada) with a 33G blunt needle was used to deliver 0.4-1 µl of the Sender AAV (1–5 × 10^12-13^ GC/ml) into the subretinal space, taking care to avoid lens damage and to achieve a localized subretinal bleb. For Receiver-AAV delivery to bipolar cells, an intravitreal injection of 0.5-1 µl of Receiver AAV at a similar titer was performed through the same incision using a similar Hamilton syringe. The animals were then placed on a heating pad (Life Brand, Toronto, ON, Canada) to fully recover from anesthesia before being returned to their cages.

For tracing of local retinal circuits in ChAT-Cre or Gad2-Cre mice, Sender AAV2/9.hSyn-DIO-(Nrx3bGFP-T2A-ChR2(H134R)-YFP), Receiver AAV2/9.hSyn-LaG17-synNotch-tTA and Reporter AAV2/9.hSyn-TRE-mRuby Reporter viruses were mixed at a 3:1:1 ratio. For Sender injections alone, hSyn-DIO-(Nrx3bGFP-T2A-ChR2(H134R) was injected undiluted, Receiver/Reporter injections included a 3:1 mix of undiluted receiver and TRE-mRuby2 diluted to 10^12^ GC/mL. When one component of the viral cocktail was excluded for controls, the remaining volume was made up with sterile PBS to preserve concentrations.

Injections via stereotaxic surgeries were performed using a Stoelting stereotaxic apparatus with a model 62 RN Hamilton syringe driven by a syringe pump (#SP3101, World Precision Instruments, Sarasota, FL, USA). Mice were induced with 4% isoflurane and maintained on 2-2.5% isoflurane throughout the operation. For testing TRACR along RGC projections, 4-5 week-old mice received simultaneous intravitreal administration of Sender-GFP AAV in retina (method as above) and stereotaxic delivery of Receiver-synNotch and Reporter AAVs to thalamus, and harvested 4-6 weeks post-injection. A 2 mm x 2 mm craniotomy was performed, and virus was injected into thalamus at AP −2.3 ML ±2.4 to 2.6 DV −2.9 to 3.0. 100 nL of virus mixture was injected at 25 nL/min to decrease viral spread. Receiver/Reporter injections included a 3:1 mix of undiluted receiver and TRE-mRuby2 (RFP) diluted to 10^12^ GC/mL. This allows for a final delivered TRE-RFP dose of 2.5 x 10^7^ viral genomes, consistent with doses previously shown to provide an optimal signal/leakiness ratio for retrograde transsynaptic tracing (Lavin et al. 2020). A dilution series of TRE Reporter virus in the presence of receiver but without sender virus confirmed the absence of sender-independent background reporter activation at this dose in the thalamus (Supp. Figure 1). For testing TRACR along the projections from secondary auditory cortex to lateral amygdala, virus was injected into the target regions using a glass micropipette connected to a Hamilton syringe via polyethylene tubing. Volumes of 1 uL of virus mixture was injected at 0.1uL/min. The micropipette remained in place for 10 minutes post-injection to minimize spread during micropipette retraction. The target regions were determined relative to bregma, secondary auditory cortex at AP −2.7 ML ±4.4 DV −3.3 and lateral amygdala at AP −1.3 ML ±3.4 DV −4.8.

### Two-photon guided electrophysiological recordings

Two photon guided electrophysiological recordings were performed as previously described (Rochon et al. 2021; Olguin et al. 2025) was used to target retinal ganglion cells in retinal explants. Briefly, mice were dark adapted for at least 2 hr, euthanized, and then retinae rapidly dissected under infrared illumination into oxygenated (95% O_2_; 5% CO_2_) Ringer’s solution. Retinae were mounted onto a filter paper (MilliporeSigma, HABG01300) with the RGC layer facing up, placed in a recording chamber, mounted on the stage of a custom-built two-photon microscope, and perfused with oxygenated Ames solution warmed to 32–34°C.

Patch electrodes (4–5 MΩ) were filled with Ringer’s solution, and fluorescein 3000 MW dextran (Thermo Scientific, D7156) was added to make the electrode visible under two-photon illumination. Signals were acquired with a MultiClamp 700B amplifier (Molecular Devices) and digitized at 20 kHz using custom software written in LabView. For spikes, the MultiClamp was put into I = 0 mode and Bessel filter set at 1 kHz. Analysis of electrophysiological signals was performed in MATLAB (Simulink) as follows. Briefly, action potentials were detected in loose patch recordings using the peakfinder function and binned (50 ms) over the entire length of the trial; firing rate histograms for each trial were then averaged and subjected to further processing based on each stimulus.

To project visual stimuli, a DLP light crafter (Texas Instruments, Dallas, TX) was used to project monochrome (410 nm) visual stimuli through a custom lens assembly that steered stimulus patterns into the back of a 20× objective (Euler et al. 2009). All visual stimuli were written in MATLAB using the psychophysics toolbox and displayed with a background intensity set to 1 × 10^4^ R*/rod/s. Moving bar stimuli consisted of a bright bar moving along its long axis in one of eight directions. The bar was 200 μm wide, 1500 μm long moving at 1000μm/s. For electrophysiological experiments, the cell-receptive field center was identified using a grid of flashing spots and a small user-controlled probe and the location with the highest response assigned as the center for all subsequent stimuli. Direction selective indices for ON, OFF, and ON-OFF RGCs were calculated using the circular variance of the cell response for all eight moving bar directions using standard methods described previously (Duan et al. 2014; Rochon et al. 2021; Olguin et al. 2025).

### Electroretinograms

Electroretinogram recordings were performed on P49 *Nrl-Cre; Elfn1^fl/fl^* animals injected with either AAV.ProA6-Elfn1-V5 or AAV.ProA6-V5. The animals were dark-adapted overnight before recording. All recordings were conducted under dim red light, using a Diagnosys Espion electroretinogram device with a ColorDome Ganzfeld stimulator and Espion software. Mice were anesthetized with an intraperitoneal injection of ketamine (100 mg/ml, Ketalean, 8KET004D, Bimeda MTC Animal Health Inc., Cambridge, ON, Canada) at 50 µg/g body weight and medetomidine (1 mg/ml, Cepetor, 236 1506 0, Modern Veterinary Therapeutics LLC, Miami, FL, USA) at 1 µg/g in sterile 0.9% NaCl (JB1300, Baxter Corp. Mississauga, ON, Canada). Body temperature was maintained at 37°C throughout the experiment using a heating pad. To prepare the eyes for recording, 1% Tropicamide (Mydriacyl, 0065-0355-03, Alcon, Mississauga, ON, Canada) eye drops were used to dilate the pupils, and topical 0.2% hypromellose gel (Genteal Tears, 0078-0429-57, Alcon, Mississauga, ON, Canada) was applied to maintain corneal hydration and electrical contact. The ground electrode was placed at the tail, the reference electrode was placed subcutaneously between the eyes, and gold loop electrodes were placed on the corneas to record. White-light (6500 K) flashes were used for light stimulation. Scotopic recordings were performed at light intensities of 0.00025, 0.001, 0.05, 0.1, 1, 2.5, 5 and 10 cd·s/m², with 10 responses recorded and averaged at an inter-stimulus interval of 5 seconds (0.2 Hz) to allow for photoreceptor recovery.

### Cortical neuron cultures

Cultures of dissociated cortices were prepared from timed pregnant females, similar to as previously described (Sengar et al. 2019). Briefly, E15 fetuses were decapitated and transferred to chilled Hank’s solution. The brain was removed and the meninges were peeled from each hemisphere. The cortices, without hippocampi, were pooled for each embryo and were mechanically dissociated using a P1000 pipette tip. The cells were plated on poly-D-lysine - coated glass coverslips in 12-well plates. The plating media was Neurobasal media (Gibco 21103-049) supplemented with fetal bovine serum (Gibco 26140), L-Glutamine (Gibco 25030-081) and B27 (Gibco 17054-044). Cultures were maintained by replacing half the media with fresh maintenance media every 3-4 days, which was Neurobasal media supplemented with L-Glutamine, B27, and 1% Penicillin-Streptomycin (Sigma-Aldrich, P4333). For AAV transfections, DIV7 neurons were incubated with AAVs diluted to 1-2 x 10^9^ GC/mL in maintenance medium for 7 days. At DIV14, neurons were fixed with 4% paraformaldehyde (PFA) + 4% sucrose for 15 minutes at room temperature.

### Tissue collection, immunohistochemistry

Brain and retina tissues were harvested 4-6 weeks after AAV injections for tracing long-range projections, and 2-4 weeks for local retinal circuits, allowing time for TRACR expression, unless otherwise noted in Results. For brain tissue collection, mice were deeply anesthetised with isoflurane and transcardially perfused with normal saline (0.9% sodium chloride) or phosphate-buffered saline (PBS) followed by 4% paraformaldehyde in phosphate-buffered saline (PBS). Brains were dissected and post-fixed in 4% paraformaldehyde at 4 degrees overnight. Brains were then embedded in 4% agarose and sectioned at 75-150 µm with a VT1000S vibratome (Leica). Free-floating sections were stored at 4°C in PBS with 0.02% sodium azide until use.

Whole-mount preparations and cryosections of retinas were prepared as described previously (Ing-Esteves et al. 2018). Eyes were removed from mice either euthanized then killed by decapitation, or transcardially perfused with PBS, and briefly fixed in ice-cold 4% paraformaldehyde. Retinas were dissected and post-fixed at 4°C for 2 hours or overnight. For some experiments, eyes were marked with a silver nitrate stick (118-395, AMG Medical Inc., Mont-Royal, Canada) on the dorsal side and then were carefully dissected from the eyecup. Retinas for wholemounts were stored in PBS with 0.02% sodium azide until use. Retinas for cryosectioning were transferred to 30% sucrose in PBS overnight, embedded in Tissue-Tek OCT (Sakura Finetek), frozen in liquid nitrogen or on chilled methylbutane (Sigma). Tissues were cryosectioned at 16-20 µm onto Superfrost Plus glass slides (Thermo Fisher Scientific, Mississauga, Canada) using a cryostat (CM3050 S, Leica Biosystems, Richmond Hill, Canada) and dried for 2 hours before storage at −20 °C with desiccant in a slide box.

For immunostainings of retina sections, slides were washed three times in PBS, then blocked and permeabilized with 2-5% donkey serum and 0.1-0.2% Triton X-100 in PBS (PBST). Floating brain sections were blocked with 5% donkey serum and 0.3% PBST. For stainings including primary antibodies raised in mouse, additional blocking of endogenous mouse IgG with was performed with AffiniPure Fab Fragment Donkey Anti-Mouse IgG (H+L) (#715-007-003, Jackson Immunoresearch) for 30 minutes at room temperature. Cryosections were probed with primary antibody in blocking buffer overnight, while floating sections were probed for three days at 4°C. After washing in PBST, samples were incubated with appropriate fluorophore-conjugated secondary antibodies in 2-5% NDS for 2-3 hours at room temperature. Where indicated, DAPI (1:5,000 from a 5 mg/mL solution, # D1306, Thermo Fisher Scientific) was included with the secondary antibodies.

For cultured neurons, fixed cells were permeabilized with 0.25% Triton X-100 for 10 minutes and blocked in 5% NDS in PBS for 1 hour. Cells were incubated with primary antibodies in blocking solution overnight at 4°C. Cell surface labelling was performed as previously described (Zhou et al. 2021). Briefly, fixed cells were blocked in 5% NDS + 1%BSA in PBS without permeabilization for 1 hour and then incubated with primary antibodies in blocking solution overnight at 4°C. Then, cells were permeabilized in 0.25% Triton X-100 for 10-15 minutes, blocked again for 1 hour in blocking solution, and incubated with primary antibodies against intracellular antigens overnight at 4°C. Cells were then incubated with secondary antibodies in blocking solution for 2 hours at room temperature, followed by DAPI (1:10,000, ThermoFisher D1306) for 10 minutes. Coverslips were mounted on microscope slides (Fisherbrand, 22-034486) with Fluoromount-G mounting medium (Southern Biotech, 0100-01).

Primary antibodies used were: Chicken anti-GFP (1:1000, Aves Labs, GFP-1010; RRID:AB_2307313); goat anti-GFP (1:500, Rockland, 600-101-215, RRID:AB_218182); rabbit anti-RFP/DsRed (1:1000, Rockland, 600-401-379; RRID:AB_2209751); chicken anti-RFP (1:500, Rockland, 600-901-379, RRID: AB_10704808); goat anti-RFP (1:1000, Rockland, 200-101-379; RRID:AB_2744552); mouse anti-myc (1:500, Millipore, 05-419; RRID:AB_309725); rabbit anti-myc (1:100-1:400, Cell Signaling Technology, 2272, RRID:AB_10692100); rat anti-HA (1:100, Roche Cat# 11867423001, RRID:AB_390918); rabbit anti-V5 (1:500, Abcam, ab206566, RRID: AB_2819156); goat anti-ChAT (1:250, Millipore, AB144P; RRID:AB_2079751); rabbit anti-SATB1 (1:100, Abcam, ab49061; RRID:AB_882454); Hoechst (1:10000, Invitrogen, H3570); mouse anti-PKC (1:500, Santa Cruz Biotechnology, Sc-8393, RRID:AB_628142); rabbit anti-Ctbp2/Ribeye (1:10000, Synaptic Systems, 193-003, RRID:AB_2086768), mouse anti-PSD95 (1:500, Abcam, AB13552, RRID:AB_300453), rabbit anti-Opn4 (1:1000, Thermo Fisher Scientific, PA1-780, RRID:AB_2267547), mouse anti-SMI132 (1:1000, Covance, SMI-32P, RRID:AB_2314912), rabbit anti-Synapsin I (1:1000, Millipore, AB1543P, RRID:AB_90757), mouse anti-Gαo (1:500, Millipore, MAB3073, RRID:AB_94671), rabbit anti-SCGN (1:500, Thermo Fisher Scientific, PA5-30393, RRID:AB_2547867); rabbit anti-SOX9 (1:500, Thermo Fisher Scientific, 702016, RRID:AB_2716879).

Secondary antibodies conjugated to Dyelight 405, Alexa Fluor 488, Alexa Fluor 568, Alexa Fluor 594, Alexa Fluor 647 or Alexa Fluor 750 (Thermo Fisher Scientific or Jackson Immunoresearch) were used at 1:500 or 1:1000. Secondary antibodies used include: Donkey anti-Goat IgG 488 (1:500, Thermo Fisher Scientific, A11055, RRID:AB_2534102); Donkey anti-Mouse IgG 488 (1:500, Thermo Fisher Scientific, RRID:AB_141607); Donkey anti-Goat IgG 488 (1:1000, Jackson ImmunoResearch Labs, 705-545-147, RRID:AB_2336933); Donkey anti-Rabbit IgG 488 (1:1000, Jackson ImmunoResearch Labs, 711-545-152, RRID:AB_2313584); Donkey anti-Goat IgG 568 (1:500, Thermo Fisher Scientific, A11057, RRID:AB_2534104); Donkey anti-Chicken IgG 555(1:500, Thermo Fisher Scientific, A78950, RRID: AB_2921072); Donkey anti-Rabbit IgG 555 (1:500-1:1000, Thermo Fisher Scientific, A31572, RRID:AB_162543); Donkey anti-Rabbit IgG 568 (1:1000, Thermo Fisher Scientific, A10042, RRID:AB_2534017); Donkey anti-Guinea Pig IgG 633 (1:500, Sigma-Aldrich, SAB4600129, RRID:AB_2890636); Donkey anti-Rabbit IgG 647 (1:500, Thermo Fisher Scientific, A-31573, RRID: AB_2536183); Donkey anti-Rabbit IgG 647 (1:1000, Jackson ImmunoResearch Labs, 711-605-152, RRID:AB_2492288); Donkey anti-Goat IgG 647 (1:1000, Thermo Fisher Scientific, A-21447, RRID:AB_141844); Donkey anti-Mouse IgG 647 (1:1000, Jackson ImmunoResearch Labs, 715-605-151, RRID:AB_2340863).

### Image Acquisition and Analyses

Immunofluorescence images were acquired using a Leica SP8, Leica Stellaris 3 or 5 (Leica Microsystems, Germany), LSM 780 or LSM810 (Carl Zeiss Inc., Thornwood, NY, USA) laser scanning confocal microscopes. Images were acquired with optimal xy- and z-resolutions defined by the objective’s numerical aperture and system settings, using 20X (NA=0.75), 40X oil (NA=1.3), or 63X glycerol (NA=1.3) lens. All the comparative images were taken using the same microscope and acquisition parameters. ImageJ/Fiji (NIH) software was used for image processing and representations, including maximal projections and brightness/contrast enhancement (http://imagej.net/Fiji/Downloads; RRID:SCR_002285) (Schindelin et al. 2012).

Laminar distributions of neurites within inner plexiform layer (IPL) were analysed using Fiji as previously described (Liu et al. 2018). Fluorescence intensity values were obtained from a line scan across the full IPL using the “Plot Profile” function. Localisation values were normalised such that the scleral border of the IPL represents 0% and the vitreal border of the IPL is indicated by 100%. Fluorescence intensity values were normalised to the minimum and maximum values within each line scan. Intensity values, in arbitrary units, were then arranged in 20 equal bins across the IPL. For retina TRACR-positive cell counts, the images were analyzed using Imaris 10 (Bitplane, Zurich, Switzerland). The spot tools in Imaris were used to count myc-Receiver, tdTomato-Reporter and marker-positive cells manually. The percentage of Reporter-positive Receiver cells or marker-positive Reporter cells was determined by counting cells from one field per section, across three sections per retina, and in at least three retinas per condition.

## Quantifications and statistical analysis

Number of experiments, animals and samples are reported in figure legends for each experiment and quantifications of TRACR cell labeling. Animals of either sex were analyzed. Statistical analyses were performed using Graph-Pad Prism software or the Real Statistics Resource Pack (Charles Zaiontz. www.real-statistics.com). All data are presented as mean ± s.e.m., unless otherwise stated. Means of two groups were compared using the two-tailed Student’s t test on condition of equivalent variances determined by the ANOVA F test, or with the Mann–Whitney nonparametric test. Means of multiple samples were compared using one-way ANOVA and Tukey’s multiple-comparisons test for pairwise analyses. To compare ERG amplitudes across multiple groups and varying light intensities, a two-way ANOVA followed by Tukey’s multiple comparisons test was used. To determine if ChAT-Cre and Gad2-Cre TRACR detected similar populations, similarity indices were calculated as previously described (Duan et al. 2014; Duan et al. 2018). Similarity indices were pooled by genotype and subjected to a one-way ANOVA to determine whether groups were significantly different. If differences were detected, posthoc pairwise tests were performed to determine the significance level reported in the figure legends. Exact p-values are reported unless the values are < 0.0001.

## ACKNOWLEDGEMENTS

We thank the following investigators for generously sharing reagents: Dr. Liang Cai for plasmid sequence encoding synNotch-RAM7; Dr. Daniel Kerschensteiner for the pAAV.Grm6S[4x]-tdTomato plasmid; Dr. Michel Cayouette for the *R26LSL-TeN*T mice, and Dr. Hongkui Zeng and the Allen Brain Institute for the Ai63(TIT-tdTomato) mice. This work was supported by: Medicine by Design Canada First Research Excellence Fund (MbDNI-2020-01) to V.A.W. and J.L.L.; Fighting Blindness Canada Research Grant to J.L.L. and A.K.; NSERC Discovery Grant (RGPIN-2023-05107) to J.L.L.; CIHR Catalyst Grant (DV2-197706) to J.L.L. and A.K.; CIHR Project Grant (PJT195688) to V.A.W. and J.L.L; NIH award EY018139 to K.A.M. J.L.L. and A.K. were supported by a Tier 2 Canada Research Chair. V.A.W is supported by Donald K. Johnson Chair in Vision Research and Tier 1 Canada Research Chair. M.T.G. was supported by a CIHR Fellowship. M.I., S.M.E. and P.P. were supported by the University of Toronto Vision Science Research Scholarship.

