## Supplemental Figures 1-6 for "TRACR: an anterograde transneuronal tracing system for genetic access across synapses and longitudinal circuit analysis"

**Supplemental Figures 1 to 6**

Lead contact: Julie L. Lefebvre

Supplementary Figure 1

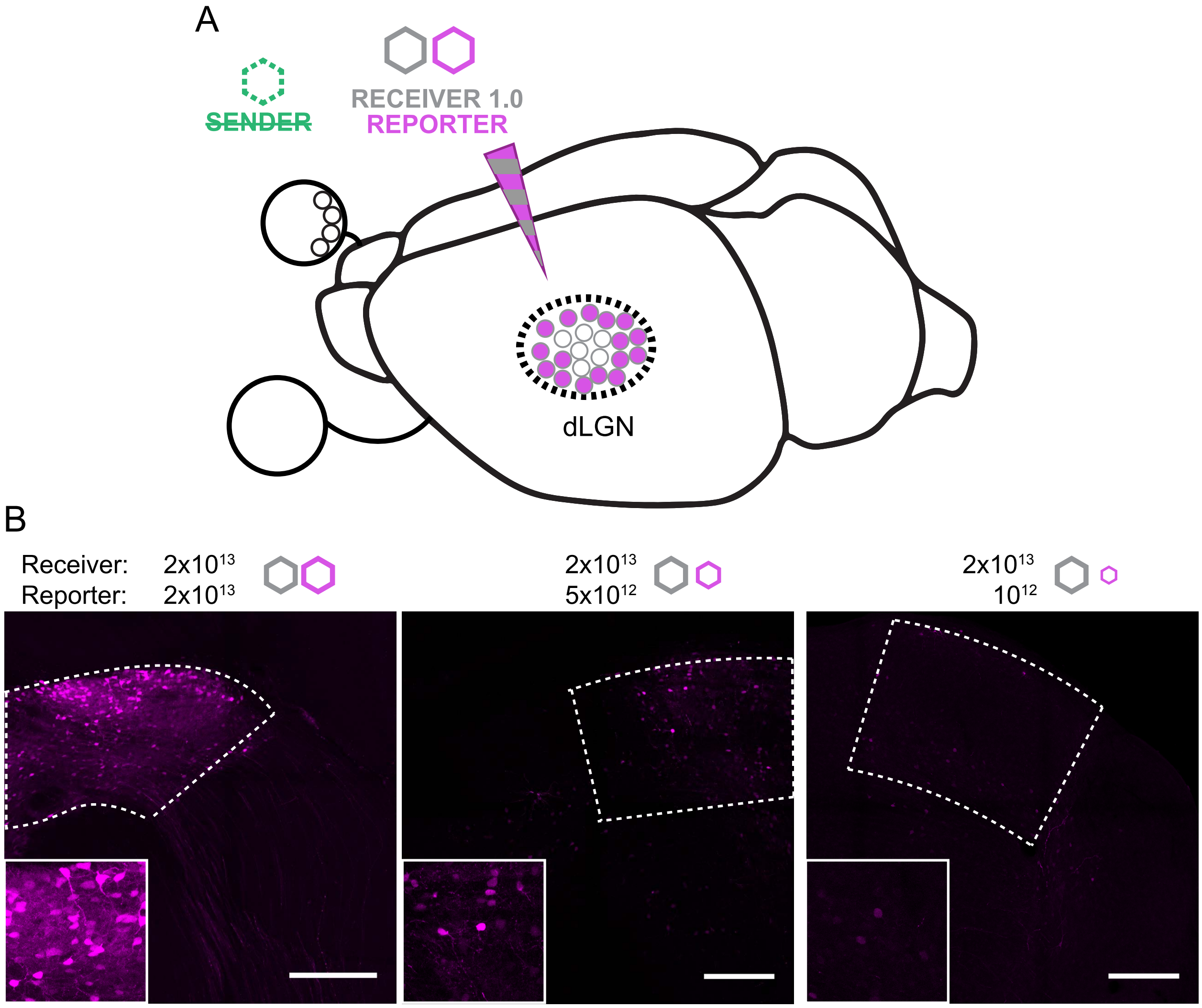


**Supplementary Figure 1. AAV control experiments.**
**(A)** To determine the appropriate reporter AAV dosage, AAVs encoding Receiver (hSyn-mycLaG17-synNotch-tTA) and Reporter (TRE-mRuby2) were injected into the thalamus in the absence of Sender AAV as uninduced controls.
**(B)** TRACR Reporter (RFP, magenta) shows significant leaky uninduced expression at high doses but relatively tight control with a 20-fold dilution. Figures are representative of 3-5 experiments per viral dose.
Scale bars: 250 μm.

Supplementary Figure 2

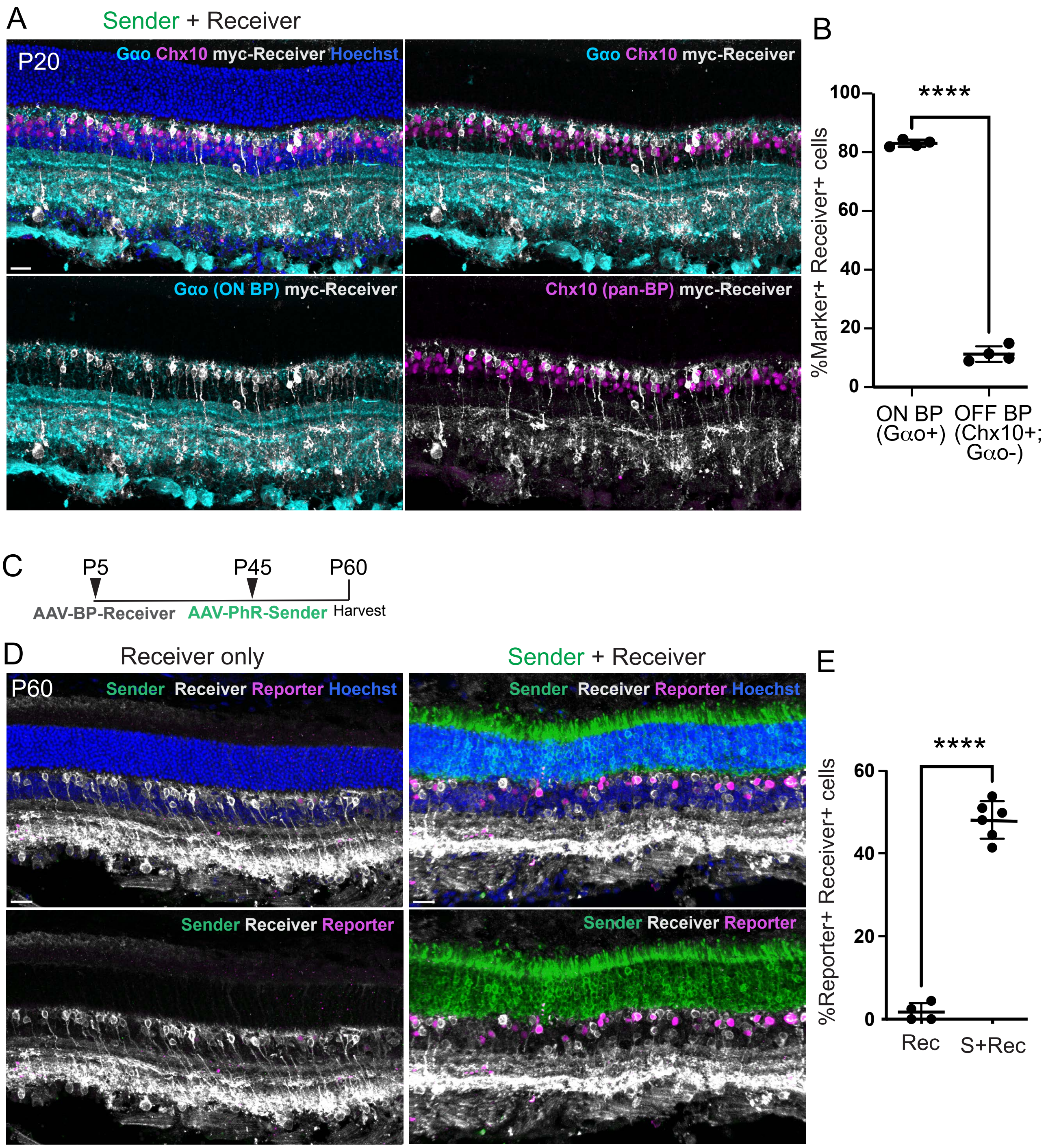

**Supplementary Figure 2. Bipolar Receivers are predominantly expressed in ON bipolar cells and the TRACR system reports established synapses in the adult retina.**
**(A)** Representative P20 retinal cross-sections from *Ai63* mice infected with the Receiver 2.0 AAV (Grm6-mycLaG17-synNotchRAM7-tTA) and stained for the ON bipolar cell marker (Goα, cyan), Receiver (myc, white), pan-bipolar cell marker (Chx10, magenta), and nuclei (Hoechst, blue). TRACR Receiver is predominantly expressed in ON bipolar cells (Chx10-positive and Go α-positive) compared with OFF bipolar cells (Chx10-positive and Goα-negative).
**(B)** Percentage of Receiver-positive cells co-expressing Chx10 and Goα. n= 3 sections per retinas, 4 retinas. Data are shown as mean ± SD. ****$p < 0.0001$.
**(C)** Schematic of AAV delivery to test TRACR labeling of established photoreceptor to bipolar synapses.  Receiver 2.0 AAV (AAV.4xGrm6-mycLaG17-synNotchRAM7-tTA) was injected into retinas of Ai63 mice at P5, followed by subretinal injection of Sender AAV (AAV.ProC1-NRXGFP-T2A-JawsHA) or PBS at P45.
**(D)** Representative P60 retinal cross-sections from *Ai63* mice infected with TRACR AAVs describe in (C). Sections were stained for Sender (GFP, green), Receiver (myc, white), tdTomato reporter (RFP, magenta), and nuclei (Hoechst, blue). **Left:** Receiver-only controls show minimal ligand-independent reporter activation. **Right:** Co-injection of Sender and Receiver AAVs results in robust Reporter activation confined to Receiver-expressing bipolar cells. (magenta).
**(E)** Percentage of Reporter-positive Receiver-expressing cells. n = 3 sections per retina and 4 retinas (Receiver only), 6 retinas (Sender + Receiver). Data are shown as mean ± SD. ****$p < 0.0001$.
Scale bars: 20 μm.

Supplementary Figure 3

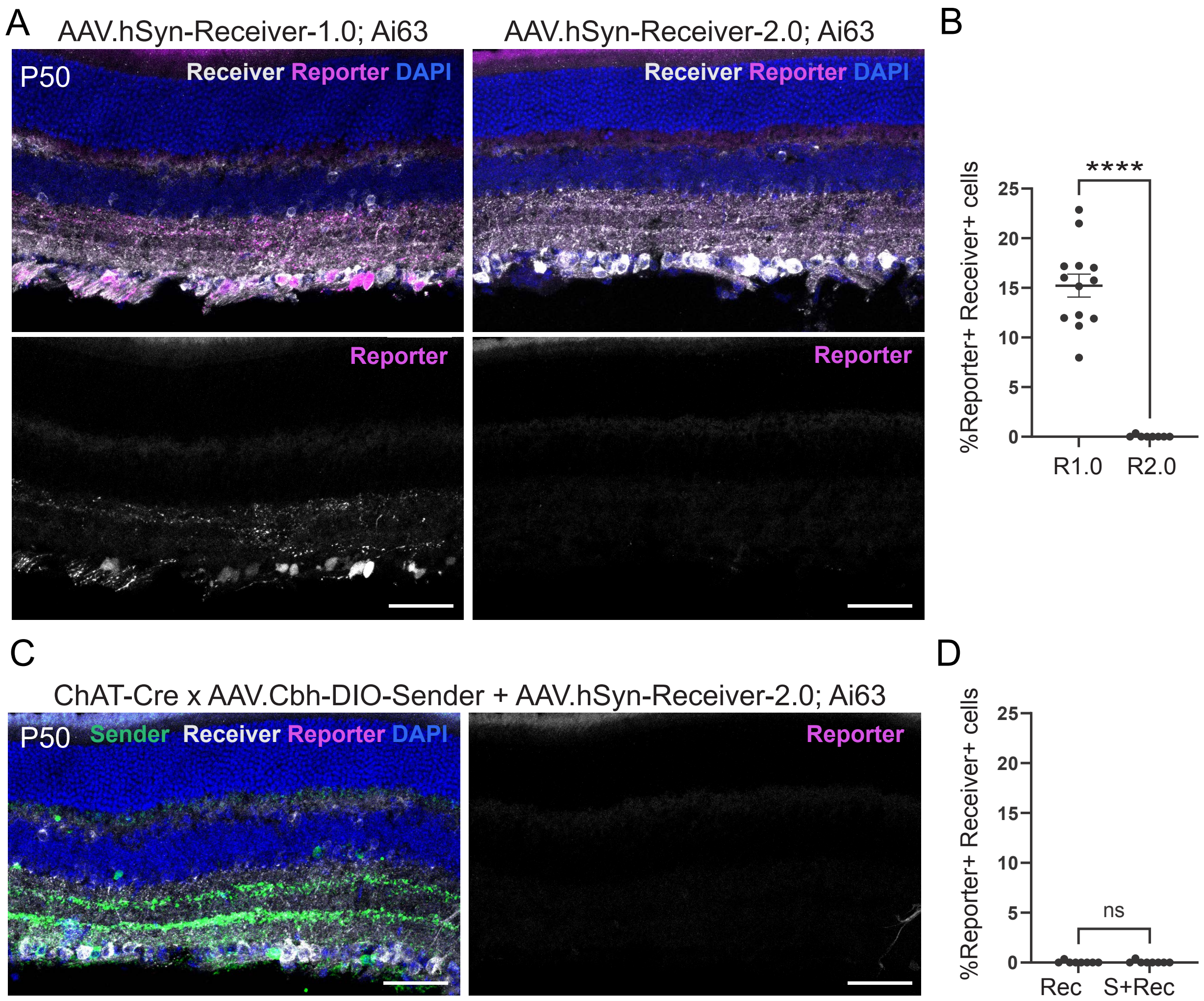


**Supplementary Figure 3. Receiver-2.0 shows minimial ligand-independent activation but fails to induce the Reporter in the presence of the Sender in *ChAT-Cre; Ai63* retinas.**

**(A)** Retinas of *ChAT-Cre; Ai63* mice were infected with AAVs encoding Receiver-1.0 (AAV7m8.hSyn-mycLaG17-synNotch-tTA) or Receiver-2.0 (AAV7m8.hSyn-mycLaG17-synNotchRAM7-tTA) at P21, and analyzed at P50. Retinal sections show Receiver (myc, white), nuclei (DAPI, blue), and Reporter (tdTomato, magenta). In the absence of Sender-GFP, ligand-independent Reporter activation is observed in Receiver-1.0 infected retinas (left) but not in Receiver-2.0 retinas (right).

**(B)** Percentage of Reporter-positive Receiver-expressing cells. N = 4 sections per retina, 13 retinas (Receiver-1.0), 8 retinas (Receiver-2.0). Data are shown as mean ± SEM. ****$p < 0.0001$, Mann-Whitney test.

**(C)** Retinal cross-sections from P50 *Chat-Cre; Ai63* mice infected with Sender (hSyn or Cbh-driven DIO-NRXGFP-T2A-ChR2-YFP) and Receiver-2.0 AAVs, and stained for Sender (GFP, green), Receiver (myc, white), and nuclei (DAPI, blue). tdTomato Reporter is unamplified (magenta). *Chat-Cre* specific Sender with pan-retinal Receiver-2.0 expression produced few Reporter-positive targets in *Chat-Cre; Ai63* mice.

**(D)** Percentage of Reporter-positive Receiver-expressing cells. n = 4 sections per retina, and 8 retinas. Data are shown as mean ± SEM. ns, $p > 0.05$, Mann-Whitney test.

Scale bars, 50µm.

Supplementary Figure 4

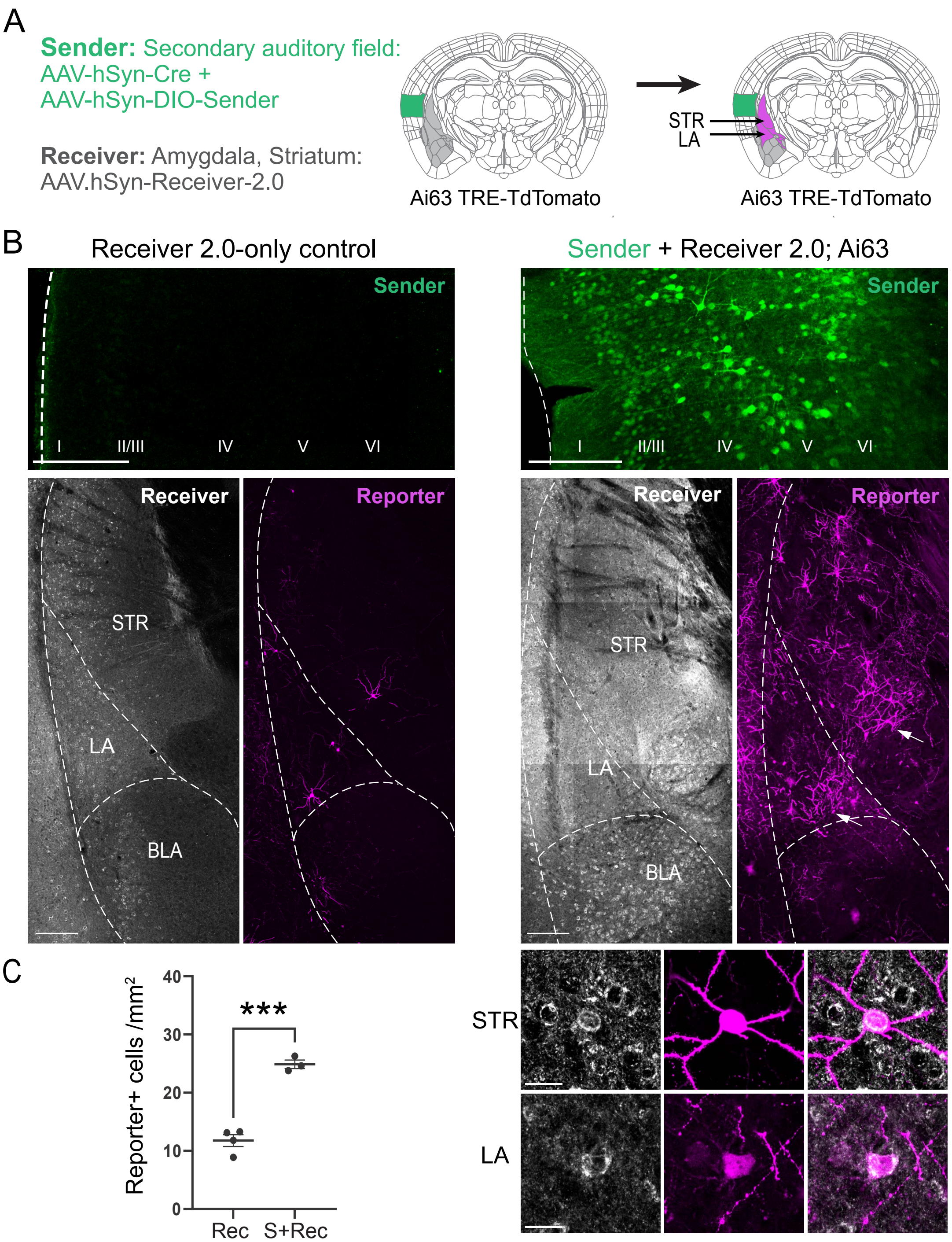

**Supplementary Figure 4. TRACR with Receiver 2.0; *Ai63* labels predicted targets of secondary auditory cortical projections in lateral amygdala and striatum.**

**(A)** Schematic of TRACR strategy to label targets of secondary auditory cortical projections. AAVs encoding the Sender (hSyn-DIO-NRXGFP-T2A-ChR2-YFP) and Cre (hSyn-Cre) were delivered by stereotaxic injection in the secondary auditory cortex (A2), and the Receiver 2.0 (hSyn-mycLaG17-synNotchRAM7-tTA) into the amygdala and surrounding striatum of *Ai63* mice. TRACR signaling (magenta) is predicted in the lateral amygdala (LA) and striatum (STR), known anatomical targets of A2 projections (LeDoux et al. 1991; Tsukano et al. 2019). Coronal brain sections were analyzed 3-4 weeks following AAV delivery, and stained for Sender (GFP, green), Receiver (myc, white), Reporter (RFP, magenta), and nuclei (DAPI, blue).

**(B) Left:** In the absence of the Sender in the A2 cortex (top), few tdTomato-positive cells (magenta) are observed in Receiver-expressing cells (white) in LA and striatal regions (STR) (bottom, dashed lines). **Right:** In brains with Sender-GFP expression in the A2 cortex, Reporter signals are observed in the Receiver-labelled LA and STR regions, compared to the Receiver-labelled basal amygdala (BLA) know to receive minor innervation from A2 (Tsukano et al. 2019). Inset shows tdTomato expression by Receiver-positive cells.

**(C)** Quantifications of tdTomato-positive cells in regions of interest spanning amygdala and striatum. n = 3 sections per brain, 4 animals (Receiver 2.0 only), 3 animals (Sender + Receiver 2.0). Data are shown as mean ± SEM. ***p = 0.0002, Student t-test.

Scale bars: 200μm in B top, 20μm in B, bottom.

Supplementary Figure 5

Muller Glia-specific Sender: ProB2-NRXGFP-T2A-JAWS$^{HA}$

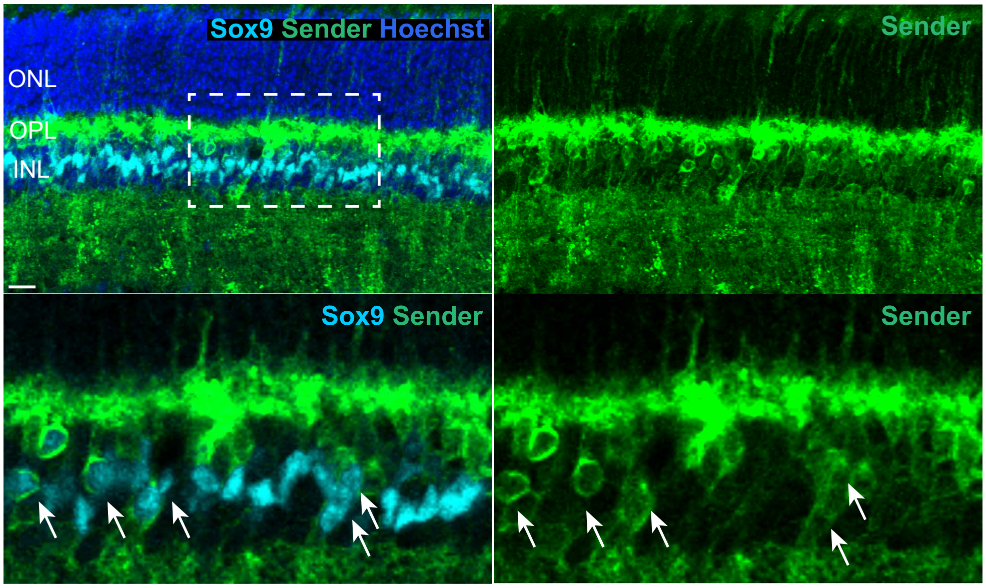


**Supplementary Figure 5. Müller glia-driven Senders are expressed in Müller glial cells.** Representative P20 retinal cross-section from *Ai63* mice infected with the Müller glial-specific AAV.MG-Sender (ProB2-NRXGFP-T2A-JAWSHA) at P5, and stained with Sender (GFP, green) and the Müller glial cell marker (SOX9, cyan). MG Sender-expressing cells show colocalization with the Müller glial marker SOX9 (white arrows).
Scale bar: 15 μm.

Supplementary Figure 6

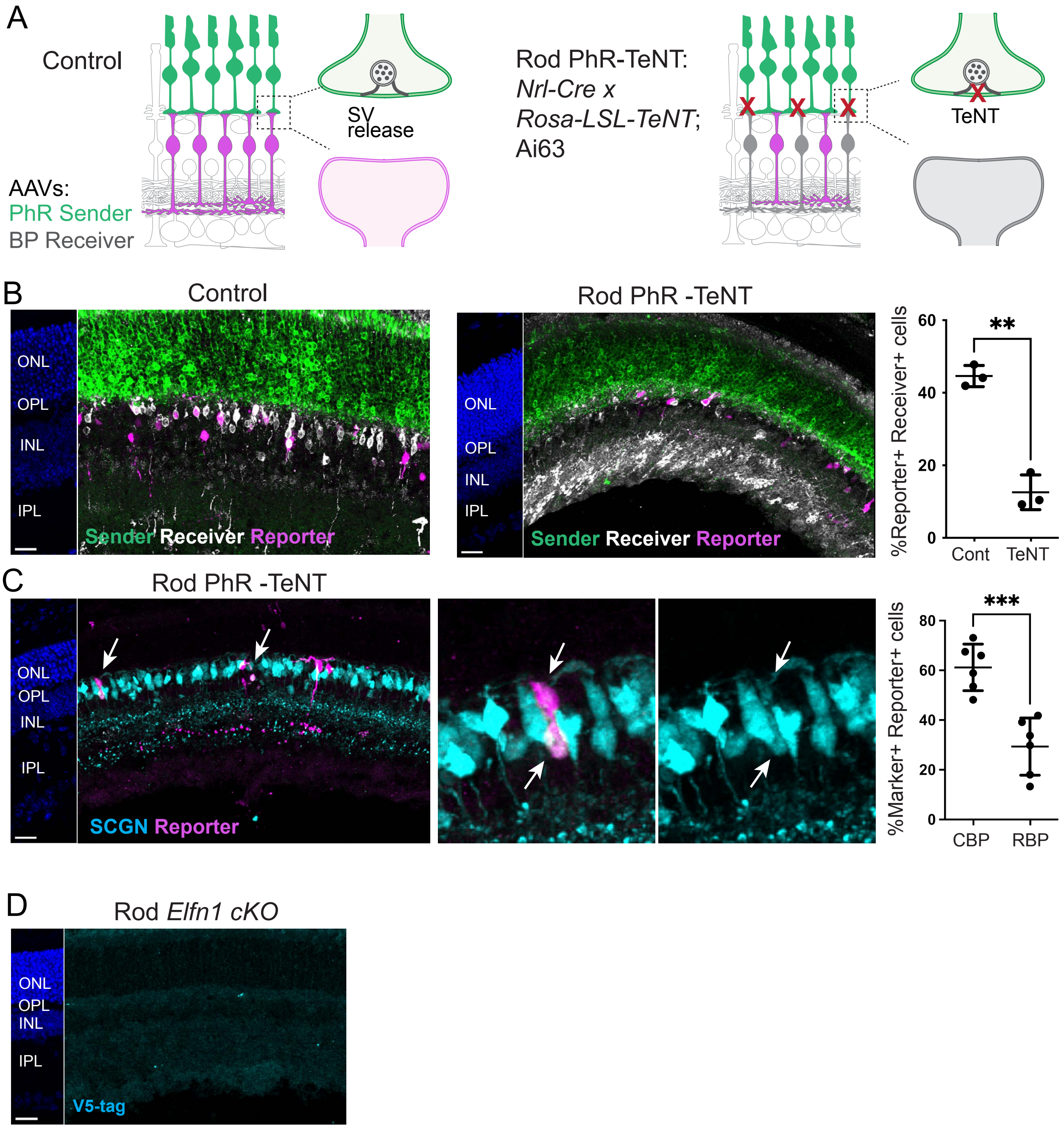

**Supplementary Figure 6. TRACR activation is reduced in an independent model of rod synapse disruption.**

**(A)** Schematic of experimental strategy to block photoreceptor neurotransmission by rod-specific expression of tetanus toxin (*Nrl-Cre;Rosa-LSL-TeNT*). Mice were crossed with the *Ai63* reporter line, infected with AAV.PhR-Sender and AAV.BP-Receiver at P5 and analyzed at P20.

**(B)** TRACR activation is reduced when neurotransmitter release from rods is blocked. **Left:** Representative P20 retinal cross-sections from control (*Rosa-TeNT;Ai63*) and *Nrl-Cre;Rosa-TeNT;Ai63* mice infected with Sender and Receiver AAVs and stained for Sender (GFP, green), Receiver (myc, white), Reporter (tdTomato, magenta), and nuclei (Hoechst, blue). **Right:** Percentage of Reporter-positive Receiver-expressing cells. n = 3 sections per retina, 3 retinas per condition. Data are shown as mean ± SD. **p = 0.0015, Welch's t-test.

**(C)** TRACR activation is preserved in cone bipolar cells when rod neurotransmitter release is blocked. **Left:** Representative retinal cross-sections from *Nrl-Cre; Rosa-TeNT;Ai63* mice stained for Reporter (tdTomato, magenta), and cone bipolar cells (SCGN, cyan). **Right:** Percentage of cone bipolar cells (SCGN-positive) or rod bipolar cells (PKC-positive) Reporter-positive cells. 3 sections per retina, 6 retinas per condition. Data are shown as mean ± SD. ***p = 0.0004, Welch's t-test.

**(D)** Control retinal sections from a *Nrl-Cre Elfn1fl/fl; Ai63* uninfected eye stained with anti-V5, confirming the specificity of V5 immunolabelling for experiments shown in Figure 6.

Scale bars: 20 µm.
